# METABOLIC: High-throughput profiling of microbial genomes for functional traits, biogeochemistry, and community-scale metabolic networks

**DOI:** 10.1101/761643

**Authors:** Zhichao Zhou, Patricia Q. Tran, Adam M. Breister, Yang Liu, Kristopher Kieft, Elise S. Cowley, Ulas Karaoz, Karthik Anantharaman

## Abstract

**Background:** Advances in microbiome science are being driven in large part due to our ability to study and infer microbial ecology from genomes reconstructed from mixed microbial communities using metagenomics and single-cell genomics. Such omics-based techniques allow us to read genomic blueprints of microorganisms, decipher their functional capacities and activities, and reconstruct their roles in biogeochemical processes. Currently available tools for analyses of genomic data can annotate and depict metabolic functions to some extent, however, no standardized approaches are currently available for the comprehensive characterization of metabolic predictions, metabolite exchanges, microbial interactions, and contributions to biogeochemical cycling.

**Results:** We present METABOLIC (**MET**abolic **A**nd **B**ioge**O**chemistry ana**L**yses **I**n mi**C**robes), a scalable software to advance microbial ecology and biogeochemistry using genomes at the resolution of individual organisms and/or microbial communities. The genome-scale workflow includes annotation of microbial genomes, motif validation of biochemically validated conserved protein residues, identification of metabolism markers, metabolic pathway analyses, and calculation of contributions to individual biogeochemical transformations and cycles. The community-scale workflow supplements genome-scale analyses with determination of genome abundance in the community, potential microbial metabolic handoffs and metabolite exchange, and calculation of microbial community contributions to biogeochemical cycles. METABOLIC can take input genomes from isolates, metagenome-assembled genomes, or from single-cell genomes. Results are presented in the form of tables for metabolism and a variety of visualizations including biogeochemical cycling potential, representation of sequential metabolic transformations, and community-scale metabolic networks using a newly defined metric ‘MN-score’ (metabolic network score). METABOLIC takes ∼3 hours with 40 CPU threads to process ∼100 genomes and metagenomic reads within which the most compute-demanding part of hmmsearch takes ∼45 mins, while it takes ∼5 hours to complete hmmsearch for ∼3600 genomes. Tests of accuracy, robustness, and consistency suggest METABOLIC provides better performance compared to other software and online servers. To highlight the utility and versatility of METABOLIC, we demonstrate its capabilities on diverse metagenomic datasets from the marine subsurface, terrestrial subsurface, meadow soil, deep sea, freshwater lakes, wastewater, and the human gut.

**Conclusion:** METABOLIC enables consistent and reproducible study of microbial community ecology and biogeochemistry using a foundation of genome-informed microbial metabolism, and will advance the integration of uncultivated organisms into metabolic and biogeochemical models. METABOLIC is written in Perl and R and is freely available at https://github.com/AnantharamanLab/METABOLIC under GPLv3.

## BACKGROUND

Metagenomics and single-cell genomics have transformed the field of microbial ecology by revealing a rich diversity of microorganisms from diverse settings, including terrestrial [1-3] and marine environments [4, 5] and the human body [6]. These approaches can provide an unbiased and insightful view into microorganisms mediating and contributing to biogeochemical activities at a number of scales ranging from individual organisms to communities [2, 7-9]. Recent studies have also enabled the recovery of hundreds to thousands of genomes from a single sample or environment [2, 8, 10, 11]. However, analyses of ever-increasing datasets remain a challenge. For example, scalable and reproducible bioinformatic approaches to characterize metabolism and biogeochemistry and standardize their analyses and representation for large datasets are lacking.

Microbially-mediated biogeochemical processes serve as important driving forces for the transformation and cycling of elements, energy, and matter among the lithosphere, atmosphere, hydrosphere, and biosphere [12]. Microbial communities in natural environmental settings exist in the form of complex and highly connected networks that share and compete for metabolites [13, 14]. The interdependent and cross-linked metabolic and biogeochemical interactions within a community can provide a relatively high level of plasticity and flexibility [2, 15]. For instance, multiple metabolic steps within a specific pathway are often separately distributed in a number of microorganisms and they are interdependent on utilizing the substrates [2, 16, 17]. This phenomenon, referred to as ‘metabolic handoffs’, is based on sequential metabolic transformations, and provides the benefit of high resilience of metabolic activities which make both the community and function stable in the face of perturbations [2, 16, 17]. It is therefore highly valuable to obtain the information of microbial metabolic function from the perspective of individual genomes as well as the entire microbial community. Our current knowledge of microbial metabolic networks is quite limited due to the lack of quantitative approaches to interpret functional details and reconstruct metabolic relationships [2]. This requires further investigation based on advanced genomic techniques and insights provided by the ever-expanding microbial genome databases.

Prediction of microbial metabolism relies on the annotation of protein function for microorganisms using a number of established databases, e.g., KEGG [18], MetaCyc [19], Pfam [20], TIGRfam [21], SEED/RAST [22], and eggNOG [23]. However, these results are often highly detailed, and therefore can be overwhelming to users. Obtaining a functional profile and identifying metabolic pathways in a microbial genome can involve manual inspection of thousands of genes [24]. Organizing, interpreting, and visualizing such datasets remains a challenge and is often untenable especially with datasets larger than one microbial genome. There is a critical need for approaches and tools to identify and validate the presence of metabolic pathways, biogeochemical function, and connections in microbial communities in a user-friendly manner. Such tools addressing this gap would also allow standardization of methods and easier integration of genome-informed metabolism into biogeochemical models, which currently rely primarily on physicochemical data and treat microorganisms as black boxes [25]. A recent statistical study indicates that incorporating microbial community structure in biogeochemical modeling could significantly increase model accuracy of processes that are mediated by narrow phylogenetic guilds via functional gene data, and processes that are mediated by facultative microorganisms via community diversity metrics [26]. This highlights the importance of integrating microbial community and genomic information into the prediction and modeling of biogeochemical processes.

Here we present the software METABOLIC, a toolkit to profile metabolic and biogeochemical functional traits based on microbial genomes. METABOLIC integrates annotation of proteins using KEGG [18], TIGRfam [21], Pfam [20], and custom hidden Markov model (HMM) databases [2], incorporates a motif validation step to accurately identify proteins based on prior biochemical validation, determines presence or absence of metabolic pathways based on KEGG modules, and produces user-friendly outputs in the form of tables and figures including a summary of functional profiles, biogeochemically-relevant pathways, and metabolic networks for individual genomes and at the community scale.

## METHODS

### HMM databases used by METABOLIC

To generate a broad range of metabolic gene HMM profiles, we integrated three sets of HMM-based databases, which are KOfam [27] (July 2019 release, containing HMM profiles for KEGG/KO with predefined score thresholds), TIGRfam [21] (Release 15.0), Pfam [20] (Release 32.0), and custom metabolic HMM profiles [2]. In order to achieve a better HMM search result excluding non-specific hits, we have tested and manually curated cutoffs for those HMM databases listed above into the resulting HMMs: KOfam database - KOfam suggested values; TIGRfam/Pfam/Custom databases - manually curated by adjusting noise cutoffs (NC) and trusted cutoffs (TC) to avoid potential false positive hits. For the KOfam suggested cutoffs, we considered both the score type (full length or domain) and the score value to assign whether an individual protein hit is significant or not. Methods on the manual curation of these databases are described in the next section.

### Curation of cutoff scores for metabolic HMMs

Two curation methods for adjusting NC or TC of TIGRfam/Pfam/Custom databases were used for a specific HMM profile. First, we parsed and downloaded representative protein sequences according to either the corresponding KEGG identifier or UniProt identifier [28]. We then randomly subsampled a small portion of the sequences (10% of the whole collection if this was more than 10 sequences, or at least 10 sequences) as the query to search against the representative protein collections [29]. Subsequently, we obtained a collection of hmmsearch scores by pair-wise sequence comparisons. We plotted scores against hmmsearch hits and selected the mean value of the sharpest decreasing interval as the adjusted cutoff. Second, we downloaded a collection of proteins that belong to a specific HMM profile and pre-checked the quality and phylogeny of these proteins by constructing and manually inspecting phylogenetic trees. We applied pre-checked protein sequences as the query search against a set of training metagenomes (data not shown). We then obtained a collection of hmmsearch scores of resulting hits from the training metagenomes. By using a similar method as described above, the cutoff was selected as the mean value of the sharpest decreasing interval.

The following example demonstrates how the method above was used to curate the hydrogenase enzymes. We then expanded this method to all genes using a similar method. We downloaded the individual protein collections for each hydrogenase functional group from the HydDB [30], which included [FeFe] Group A-C series, [Fe] Group, and [NiFe] Group 1-4 series. The individual hydrogenase functional groups were further categorized based on the catalyzing directions, which included H2-evolution, H2-uptake, H2-sensing, electron-bifurcation, and bidirection. To define the NC cutoff (‘--cut_nc’ in hmmsearch) for individual hydrogenase groups, we used the protein sequences from each hydrogenase group as the query to hmmsearch against the overall hydrogenase collections. By plotting the resulting hmmsearch hit scores against individual hmmsearch hits, we selected the mean value of the sharpest decreasing interval as the cutoff value.

### Motif validation

To automatically validate protein hits and avoid false positives, we introduced a motif validation step by comparing protein motifs against a manually curated set of highly conserved residues in important proteins. This manually curated set of highly conserved residues is derived from either reported works or protein alignments from this study. We chose 20 proteins associated with important metabolisms (with a focus on important biogeochemical cycling steps) that are prone to being misannotated into proteins within the same protein family. Details of these proteins are provided in Additional file 8: Dataset S1. For example, DsrC (sulfite reductase subunit C) and TusE (tRNA 2-thiouridine synthesizing protein E) are similar proteins that are commonly misannotated. Both of them are assigned to the family KO:K11179 in the KEGG database. To avoid assigning TusE as a sulfite reductase, we identified a specific motif for DsrC but not TusE (GPXKXXCXXXGXPXPXXCX”, where “X” stands for any amino acid) [31]. We used these specific motifs to filter out proteins that have high sequence similarity but functionally divergent homologs.

### Annotation of carbohydrate-active enzymes and peptidases

For carbohydrate-active enzymes (CAZymes), dbCAN2 [32] was used to annotate proteins with default settings. The hmmscan parser and HMM database (2019-09-05 release) were downloaded from the dbCAN2 online repository (http://bcb.unl.edu/dbCAN2/download/) [32]. The non-redundant library of protein sequences which contains all the peptidase/inhibitor units from the peptidase (inhibitor) database MEROPS [33] was used as the reference database to search against putative peptidases and inhibitors using DIAMOND. The settings used for the DIAMOND BLASTP search were “-k 1 -e 1e-10 --query-cover 80 --id 50” [34]. We used the ‘MEROPS pepunit’ database since it only includes the functional unit of peptidases/inhibitors [33] which can effectively avoid potential non-specific hits.

### Implementation of METABOLIC-G and METABOLIC-C

To target specific applications in processing omics datasets, we have implemented two versions of METABOLIC – METABOLIC-G (genome version) and METABOLIC-C (community version). METABOLIC-G intakes only genome files and provides analyses for individual genome sequences. METABOLIC-C includes an option for users to include metagenomic reads for mapping to metagenome-assembled genomes (MAGs).

Using Bowtie 2 (version ≥ v2.3.4.1) [35], metagenomic bam files were generated by mapping all input metagenomic reads to gene collections from input genomes. Subsequently, SAMtools (version ≥ v0.1.19) [36], BAMtools (version ≥ v2.4.0) [37], and CoverM (https://github.com/wwood/CoverM) were used to convert bam files to sorted bam files and to calculate the gene depth of read coverage. To calculate the relative abundance of a specific biogeochemical cycling step, all the coverage of genes that are responsible for this step were summed up and normalized by overall gene coverage. Reads from single-cell and isolate genomes can also be mapped in an identical manner to metagenomes. The gene coverage result generated by metagenomic read mapping was further used in downstream processing steps to conduct community-scale interaction and network analyses.

### Classifying microbial genomes into taxonomic groups

To study community-scale interactions and networks of each microbial group within the whole community, we classified microbial genomes into individual taxonomic groups. GTDB-Tk v0.1.3 [38] was used to assign taxonomy of input genomes with default settings. GTDB-Tk can provide automated and objective taxonomic classification based on the rank-normalized Genome Taxonomy Database (GTDB) taxonomy within which the taxonomy ranks were established by a sophisticated criterion counting the relative evolutionary divergence (RED) and average nucleotide identity (ANI) [38, 39]. Subsequently, genomes were clustered into microbial groups at the phylum level, except for Proteobacteria which were replaced by its subordinate classes due to its wide coverage. Taxonomic assignment information for each genome was used in the downstream community analyses.

### Analyses and visualization of metabolic outputs, biogeochemical cycles, MN-scores, metabolic networks, and energy flow potential

To visualize the outputted metabolic results, R script “*draw_biogeochemical_cycles*.*R*” was used to draw the corresponding metabolic pathways for individual genomes. We integrated HMM profiles that are related to biogeochemical activities and assigned HMM profiles to 31 distinct biogeochemical cycling steps (See details in “METABOLIC_template_and_database” folder on the GitHub page). The script can generate figures showing biogeochemical cycles for individual genomes and the summarized biogeochemical cycle for the whole community. By using the results of metabolic profiling generated from HMM search and gene coverage from the mapping of metagenomic reads, we can depict metabolic capacities of both individual genomes and all genomes within a community as a whole. The community-level diagrams, including sequential transformations, metabolic energy flow, and metabolic network diagrams, were generated using both metabolic profiling and gene coverage results. The diagrams are made by the scripts “*draw_sequential_reaction*.*R*” (using R package “*ggplot2*” [40]), “*draw_metabolic_energy_flow*.*R*” (using R package “*ggalluvial*” [41]), and “*draw_metabolic_network*.*R*” (using R package “*ggraph*” [42]), respectively (For details, refer to GitHub README page).

MN-score (metabolic network score) is a metric reflecting the functional capacity and abundance of a microbial community in co-sharing metabolic networks. It was calculated at the community-scale level based on results of metabolic profiling and gene coverage from metagenomic read mapping as described above. Metabolic potential for the whole community was profiled into individual functions that either mediated specific pathways or transformed certain substrates into products; MN-score for each function indicates its distribution weight within the metabolic networks which was calculated by summing up all the coverage values of genes belonging to the function and subsequently normalizing it by overall gene coverage. For each function, the contribution percentage of each microbial phylum in the microbial community was also calculated accordingly. Detailed description for calculating MN-scores are further provided in the results section.

### Example of metabolic diagrams

An example of community-scale analyses including element biogeochemical cycling and sequential reaction analyses, metabolic network and energy flow potential analyses, and MN-score calculation were conducted using a metagenomic dataset of microbial community inhabiting deep-sea hydrothermal vent environment of Guaymas Basin in the Pacific Ocean [43]. It contains 98 MAGs and 1 set of metagenomic reads (genomes were available at NCBI BioProject PRJNA522654 and metagenomic reads were deposited to NCBI SRA with accession as SRR3577362).

A recent metagenomic-based study of the microbial community from an aquifer adjacent to Colorado River, located near Rifle, has provided an accurate reconstruction of the metabolism and ecological roles of the microbial majority [2]. From underground water and sediments of the terrestrial subsurface at Rifle, 2545 reconstructed MAGs were obtained (genomes are under NCBI BioProject PRJNA288027). They were used as the *in silico* dataset to test METABOLIC’s performance. First, all the microbial genomes were dereplicated by dRep v2.0.5 [44] to pick the representative genomes for downstream analysis using the setting of ‘-comp 85’. Then, METABOLIC-G was applied to profile the functional traits of these representative genomes using default settings. Finally, the metabolic profile chart was depicted by assigning functional traits to GTDB taxonomy-clustered genome groups.

### Test of software performance for different environments

To benchmark and test the performance of METABOLIC in different environments, eight datasets of metagenomes and metagenomic reads from marine, terrestrial, and human environments were used. These included marine subsurface sediments [45] (Deep biosphere beneath Hydrate Ridge offshore Oregon), freshwater lake [46] (Lake Tanganyika, eastern Africa), colorectal cancer (CRC) patient gut [47], healthy human gut [47], deep-sea hydrothermal vent (Guaymas Basin, Gulf of California) [43], terrestrial subsurface sediments and water (Rifle, CO, USA) [2], meadow soils [48] (Angelo Coastal Range Reserve, CA, USA), and advanced water treatment facility [49] (Groundwater Replenishment System, Orange County, CA, USA). Default settings were used for running METABOLIC-C.

### Comparison of community-scale metabolism

To compare the metabolic profile of two environments at the community scale, MN-score was used as the benchmarker. Two sets of environment pairs were compared, including marine subsurface sediments [45] and terrestrial subsurface sediments and water [2], and freshwater lake [46] and deep-sea hydrothermal vent [43]. To demonstrate differences between these environments to specific biogeochemical processes, we focused on the biogeochemical cycling of sulfur. The sulfur biogeochemical cycling diagrams were depicted according to the number of genomes and genome coverage of organisms that contain each biogeochemical cycling step.

### Metabolism in human microbiomes

To inspect the metabolism of microorganisms in the human microbiome (associated with skin, oral mucosa, conjunctiva, gastrointestinal tracts, etc.), a subset of KOfam HMMs (139 HMM profiles) were used as markers to depict the human microbiome metabolism (parsed by HuMiChip targeted functional gene families [50]). They included 10 function categories as follows: amino acid metabolism, carbohydrate metabolism, energy metabolism, glycan biosynthesis and metabolism, lipid metabolism, metabolism of cofactors and vitamins, metabolism of other amino acids, metabolism of terpenoids and polyketides, nucleotide metabolism, and translation. The CRC and healthy human gut (healthy control) sample datasets were used as the input (Accession IDs: Bioproject PRJEB7774 Sample 31874 and Sample 532796). Heatmap of presence/absence of these functions were depicted by R package “*pheatmap*” [51] with 189 horizontal entries (there are duplications of HMM profiles among function categories; for detailed human microbiome metabolism markers refer to Additional file 9: Dataset S2).

### Representation of microbial cell metabolism

To provide a schematic representation of the metabolism of microbial cells, two microbial genomes were used as examples, Hadesarchaea archaeon 1244-C3-H4-B1 and Nitrospirae bacteria M_DeepCast_50m_m2_151. METABOLIC-G results of these two genomes, including functional traits and KEGG modules, were used to draw the cell metabolism diagrams.

### Metatranscriptome analysis by METABOLIC

METABOLIC-C can take metatranscriptomic reads as input into transcript coverage calculation and integrate the result to downstream community analyses. METABOLIC-C uses the same method as that of gene coverage calculation, including mapping transcriptomic reads to the gene collection from input genomes, converting bam files to sorted bam files, and calculating the transcript coverage. The raw transcript coverage was further normalized by the gene length and metatranscriptomic read number in Reads Per Kilobase of transcript, per Million mapped reads (RPKM). Hydrothermal vent and background seawater transcriptomic reads from Guaymas Basin (NCBI SRA accessions SRR452448 and SRR453184) were used to test the outcome of metatranscriptome analysis.

## RESULTS AND DISCUSSION

Given the ever-increasing number of microbial genomes from microbiome studies, we developed METABOLIC to enable scalable analyses of metabolic pathways and enable visualization of biogeochemical cycles and community-scale metabolic networks. METABOLIC is the first software to elucidate community-scale networks of metabolic tradeoffs, energy flow, and metabolic connections based on genome composition. While METABOLIC relies on microbial genomes and metagenomic reads for underpinning its analyses, it can easily integrate transcriptomic datasets to provide an activity-based measure of community networks.

### Workflow to determine the presence of metabolic pathways in microbial genomes

METABOLIC is written in Perl and R and is expected to run on Unix, Linux, or macOS. The prerequisites are described on METABOLIC’s GitHub page (https://github.com/AnantharamanLab/METABOLIC). The input folder requires microbial genome sequences in FASTA format and an optional set of genomic/metagenomic reads which were used to reconstruct those genomes (Figure 1). Genomic sequences are annotated by Prodigal [52], or a user can provide self-annotated proteins (with extensions of “.faa”) to facilitate incorporation into existing pipelines. We have also included an accessory Perl script which can help users to parse out the gene and protein sequences out of input genomes based on the Prodigal-generated “.gff” files. These files are used in the downstream steps involving the mapping of genomic/metagenomic reads.

**Figure 1.**
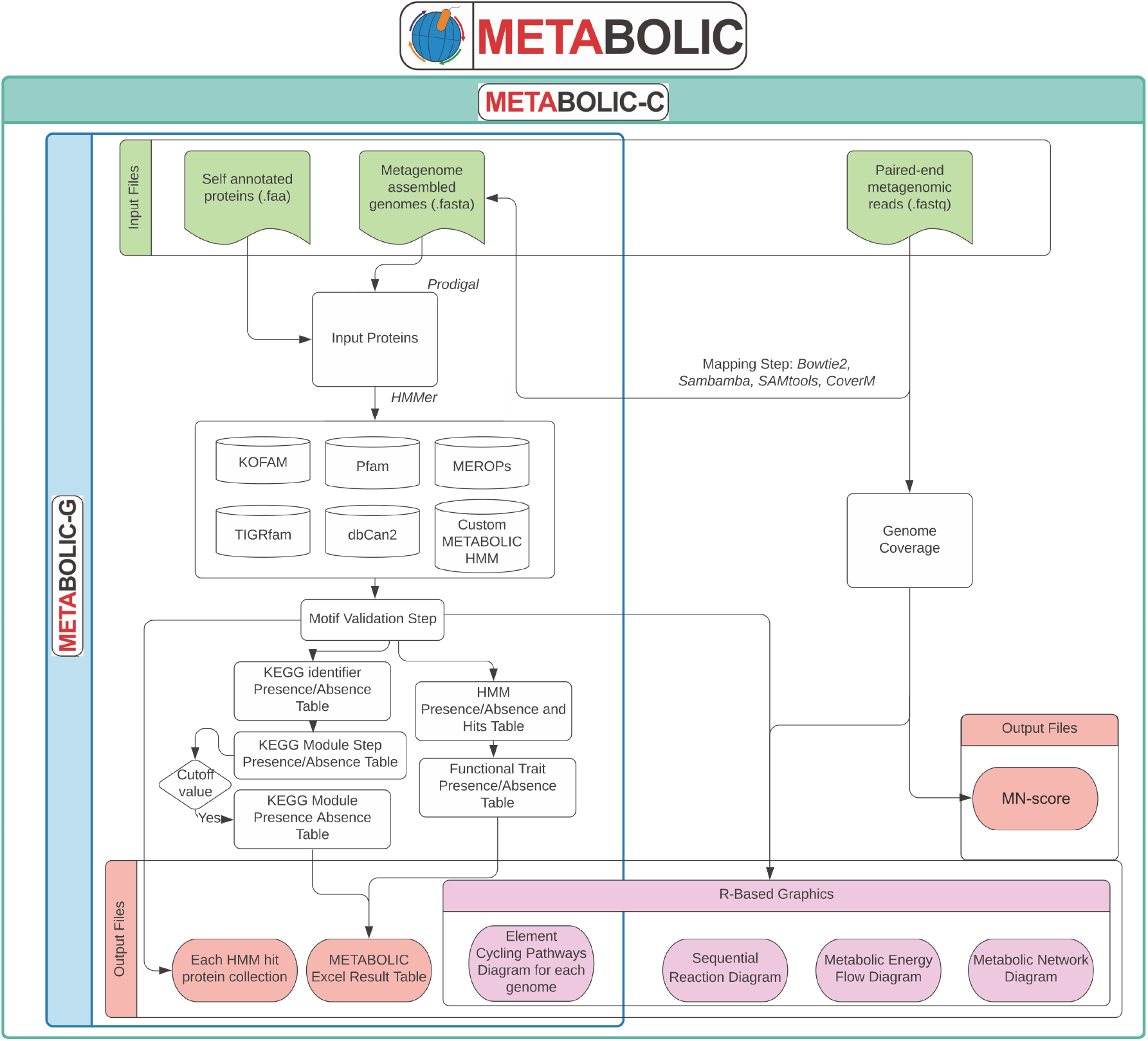
An outline of the workflow of METABOLIC. Detailed instructions are available at https://github.com/AnantharamanLab/METABOLIC. METABOLIC-G workflow was specifically shown in the blue square and METABOLC-C workflow was shown in the green square.

Proteins are queried against HMM databases (KEGG KOfam, Pfam, TIGRfam, and custom HMMs) using hmmsearch implemented within HMMER [29] which applies methods to detect remote homologs as sensitively and efficiently as possible. After the hmmsearch step, METABOLIC subsequently validates the primary outputs by a motif-checking step for a subset of protein families; only those protein hits which successfully pass this step are regarded as significant hits.

METABOLIC relies on matches to the above databases to infer the presence of specific metabolic pathways in microbial genomes. Individual KEGG annotations are inferred in the context of KEGG modules for a better interpretation of metabolic pathways. A KEGG module is comprised of multiple steps with each step representing a distinct metabolic function. We parsed the KEGG module database [53] to link the existing relationship of KO identifiers to KEGG module identifiers to project our KEGG annotation result into the metabolic network which was constructed by individual building blocks – modules – for better representation of metabolic blueprints of input genomes. In most cases, we used KOfam HMM profiles for KEGG module assignments. For a specific set of important metabolic marker proteins and commonly misannotated proteins, we also applied the TIGRfam/Pfam/custom HMM profiles and motif-validation steps. The software has customizable settings for increasing or decreasing the priority of specific databases, primarily meant to increase annotation confidence by preferentially using custom HMM databases over KEGG KOfam when targeting the same set of proteins.

Since individual genomes from metagenomes and single-cell genomes can often have incomplete metabolic pathways, we provide an option to determine the completeness of a metabolic pathway (or a module here). A user-defined cutoff is used to estimate the completeness of a given module (the default cutoff is the presence of 75% of metabolic steps/genes within a given module), which is then used to produce a KEGG module presence/absence table. All modules exceeding the cutoff are determined to be complete. Meanwhile, the presence/absence information for each module step is also summarized in an overall output table to facilitate further detailed investigations.

Outputs consist of six different results that are reported in an Excel spreadsheet (Additional file 1: Figure S1). These contain details of protein hits (Additional file 1: Figure S1A) which include both presence/absence and protein names, presence/absence of functional traits (Additional file 1: Figure S1B), presence/absence of KEGG modules (Additional file 1: Figure S1C), presence/absence of KEGG module steps (Additional file 1: Figure S1D), CAZyme hits (Additional file 1: Figure S1E) and peptidase/inhibitor hits (Additional file 1: Figure S1F). For each HMM profile, the protein hits from all input genomes can be used for the construction of phylogenetic trees or further be combined with additional datasets or reference protein collections for detailed evolutionary analyses.

### Elemental cycling pathway analyses enable quantitative calculation of microbial contributions to biogeochemical cycles

The software identifies and highlights specific pathways of importance in microbiomes associated with energy metabolism and biogeochemistry. To visualize pathways of biogeochemical importance, the software generates schematic profiles for nitrogen, carbon, sulfur, and other elemental cycles for each genome. The set of genomes used as input is considered the “community”, and each genome within is considered an “organism” when doing these calculations. A summary schematic diagram at the community level integrates results from all individual genomes within a given dataset (Figure 2) and includes computed abundances for each step in a biogeochemical cycle if the genomic/metagenomic read datasets are provided. The genome number labeled in the figure indicates the number/quantity of genomes that contain the specific gene components of a biogeochemical cycling step (Figure 2) [2]. In other words, it represents the number of organisms within a given community inferred to be able to perform a given metabolic or biogeochemical transformation. The abundance percentage indicates the relative abundance of microbial genomes that contain the specific gene components of a biogeochemical cycling step among all microbial genomes in a given community (Figure 2) [2].

**Figure 2.**
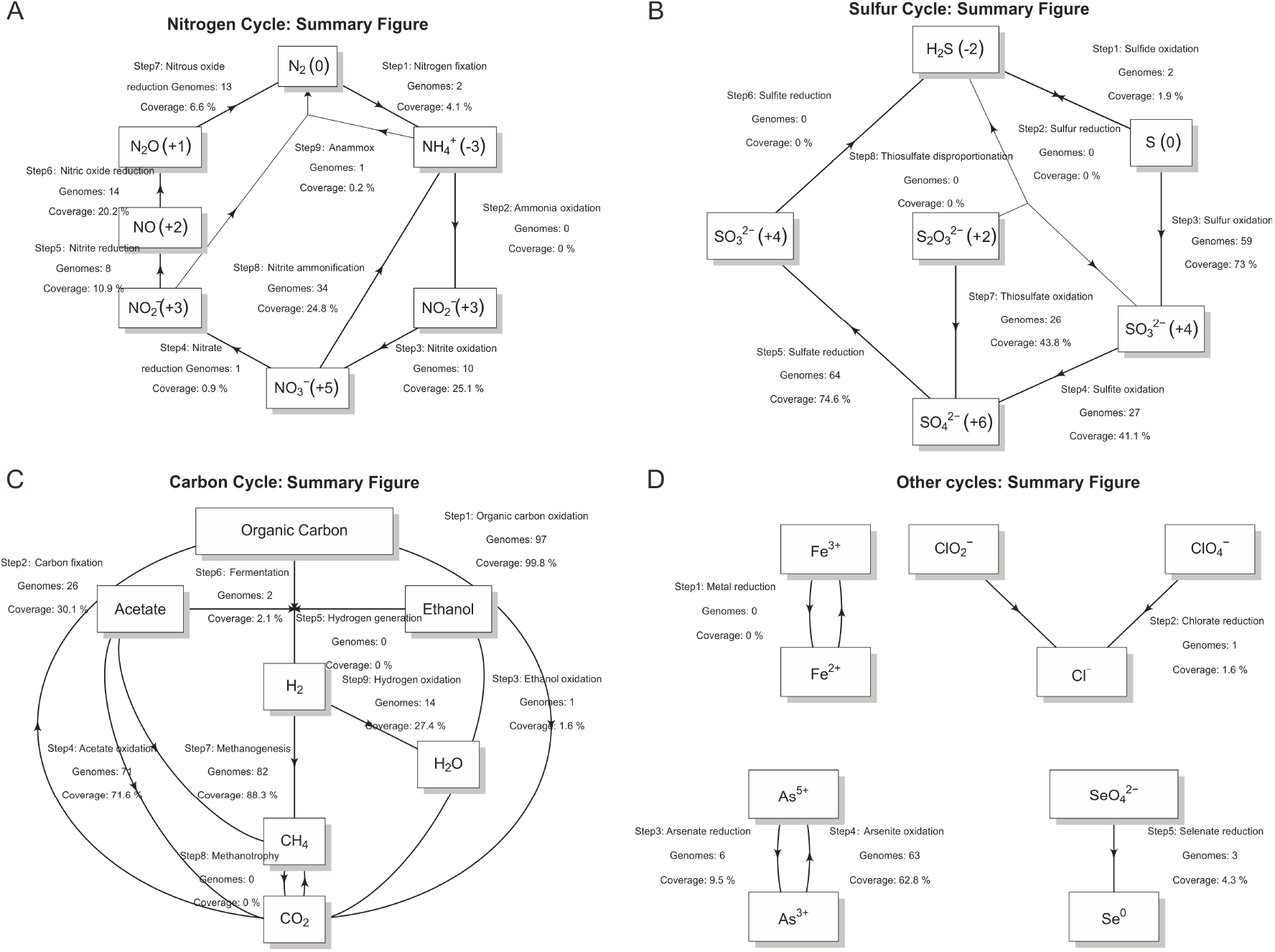
Summary scheme of biogeochemical cycling processes at the community scale. Each arrow represents a single transformation/step within a cycle. Labels above each arrow are (from top to bottom): step number and reaction, number of genomes that can conduct these reactions, metagenomic coverage of genomes (represented as a percentage within the community) that can conduct these reactions.

### Elucidating sequential reactions involving inorganic and organic compounds

Microorganisms in nature often do not encode pathways for the complete transformation of compounds. For example, microorganisms possess partial pathways for denitrification that can release intermediate compounds like nitrite, nitric oxide, and nitrous oxide in lieu of nitrogen gas which is produced by complete denitrification [54]. A greater energy yield could be achieved if one microorganism conducts all steps associated with a pathway (such as denitrification) [2] since it could fully use all available energy from the reaction. However, in reality, few organisms in microbial communities carry out multiple steps in complex pathways; organisms commonly rely on other members of microbial communities to conduct sequential reactions in pathways [2, 55, 56]. METABOLIC summarizes and enables visualization of the genome number and coverage (relative abundance) of microorganisms that are putatively involved in the sequential transformation of both important inorganic and organic compounds (Figure 3). This provides a qualitative and quantitative calculation of microbial interactions and connections using shared metabolites associated with inorganic and organic transformations.

**Figure 3.**
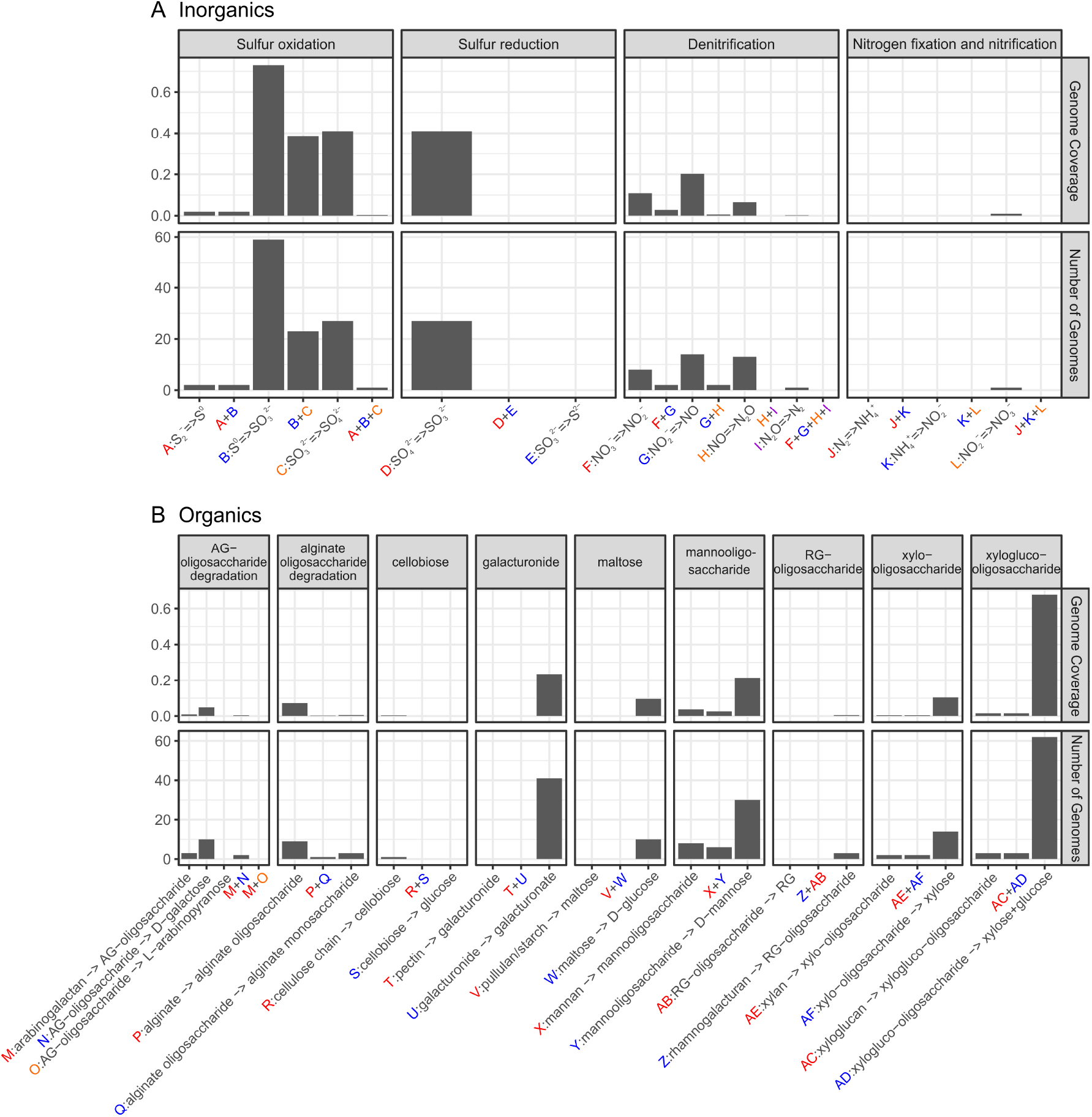
Schematic figure of sequential metabolic transformations. **(A)** the sequential transformation of inorganic compounds; **(B)** the sequential transformation of organic compounds. X-axes describe individual sequential transformations indicated by letters. The two panels describe the number of genomes and genome coverage (represented as a percentage within the community) of organisms that are involved in certain sequential metabolic transformations. The deep-sea hydrothermal vent dataset was used for these analyses.

### Construction of metabolic networks to infer connections between microbial metabolism and biogeochemical cycles

Given the abundance of microbial pathway information generated by METABOLIC, we identified co-existing metabolisms in microbial genomes as a measure of connections between different metabolic functions and biogeochemical steps. In the context of biogeochemistry, this approach allows the evaluation of relatedness among biogeochemical steps and the connection contribution by microorganisms. This is enabled at the resolution of individual genomes using the phylogenetic classification (Figure 4) assigned by GTDB-tk [38]. As an example, we have demonstrated this approach on a microbial community inhabiting deep-sea hydrothermal vents. We divided the microbial community of deep-sea hydrothermal vents into 18 phylum-level groups (except for Proteobacteria which were divided into their subordinate classes). The metabolic connection network diagrams were depicted at the resolution of both individual phyla and the entire community level (Additional file 10: Dataset S3). Figure 4 demonstrates metabolic connections that were represented with individual metabolic/biogeochemical cycling steps depicted as nodes, and the connections between two given nodes depicted as edges. The size of a given node is proportional to the gene coverage associated with the metabolic/biogeochemical cycling step. The thickness of a given edge was depicted based on the average of gene coverage values of these two biogeochemical cycling steps (the connected nodes). More edges connecting two nodes represent more connections between these two steps. The thickness of edges represents gene coverages (measured as the average of these two steps). The color of the edge corresponds to the taxonomic group, and at the whole community level, more abundant microbial groups were more represented in the diagram (Figure 4). Overall, METABOLIC provides a comprehensive approach to construct and visualize metabolic networks associated with important pathways in energy metabolism and biogeochemical cycling in microbial communities and ecosystems.

**Figure 4.**
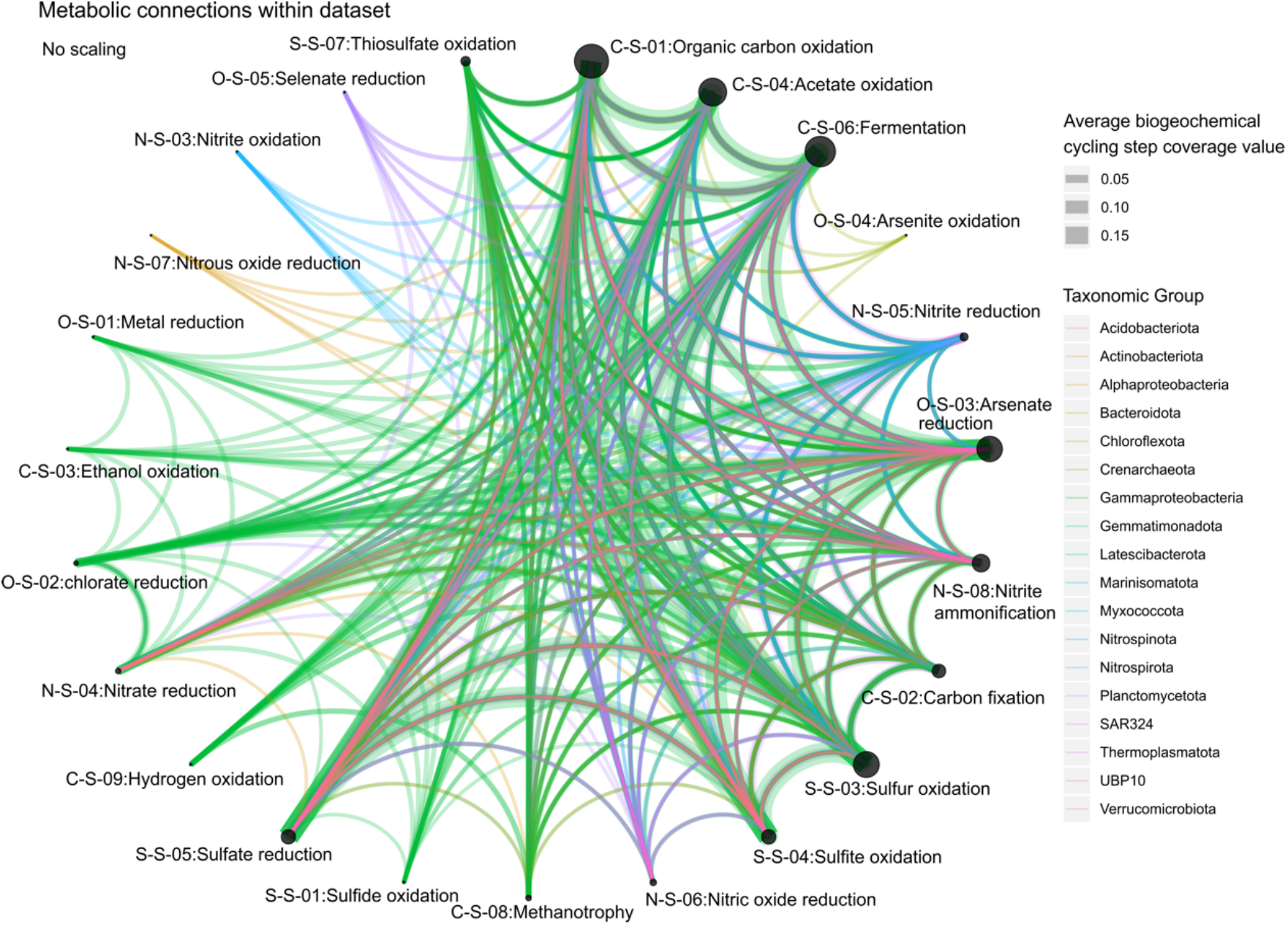
Metabolic network showing connections between different metabolisms in the microbial community. Nodes represent individual steps in biogeochemical cycles; edges connecting two given nodes represent the metabolic connections between nodes, which is enabled by organisms that can conduct both biogeochemical processes/steps. The thickness of the edge was depicted according to the average of gene coverage values of the two connected biogeochemical cycling steps – for example, thiosulfate oxidation and organic carbon oxidation.. The color of the edges was assigned based on the taxonomy of the represented genome. The deep-sea hydrothermal vent dataset was used for these analyses.

### Calculating MN-scores to represent function weights and microbial group contribution in metabolic networks

To address the lack of quantitative and reproducible measures to represent potential metabolic exchange and interactions in microbial communities, we developed a new metric that we termed MN-score (metabolic networking scores). MN-scores quantitatively measure “function weights” within a microbial community as reflected by the metabolic profile and gene coverage. As metabolic potential for the whole community was profiled into individual functions that either mediated specific pathways or transformed certain substrates into products, a function weight that reflects the abundance fraction for each function can be used to represent the overall metabolic potential of the community. MN-scores resolved the functional capacity and abundance in the co-sharing metabolic networks as studied and visualized in the above section. Towards this (Figure 5), we divided metabolic/biogeochemical cycling steps (31 in total) into a finer level – function (51 functions in total) – for better resolution on reflecting metabolic networks. By using similar methods for determining metabolic interactions (as in the above section), we selected functions that are shared among genomes and summarized their weights within the whole community by adding up their abundances. More frequently shared functions and their higher abundances lead to higher MN-scores, which quantitively reflects the function weights in metabolic networks (Figure 5). MN-score reflects the same metabolic networking pattern with the above description on the edges (networking lines) connecting the nodes (metabolic steps) that – more edges connecting two nodes indicates two steps are more co-shared, thicker edges indicate higher gene abundance for the metabolic steps. The MN-scores can integratively represent these two networking patterns and serve as metrics to measure these function weights. At the same time, we also calculated each microbial group’s (phylum in this case) contribution to the MN-score of a specific function within the community (Figure 5). A higher microbial group contribution percentage value indicates that one function is more represented by the microbial group (for both gene presence and abundance) in the metabolic networks. MN-scores provide a quantitive measure on comparing function weights and microbial group contributions within metabolic networks.

**Figure 5.**
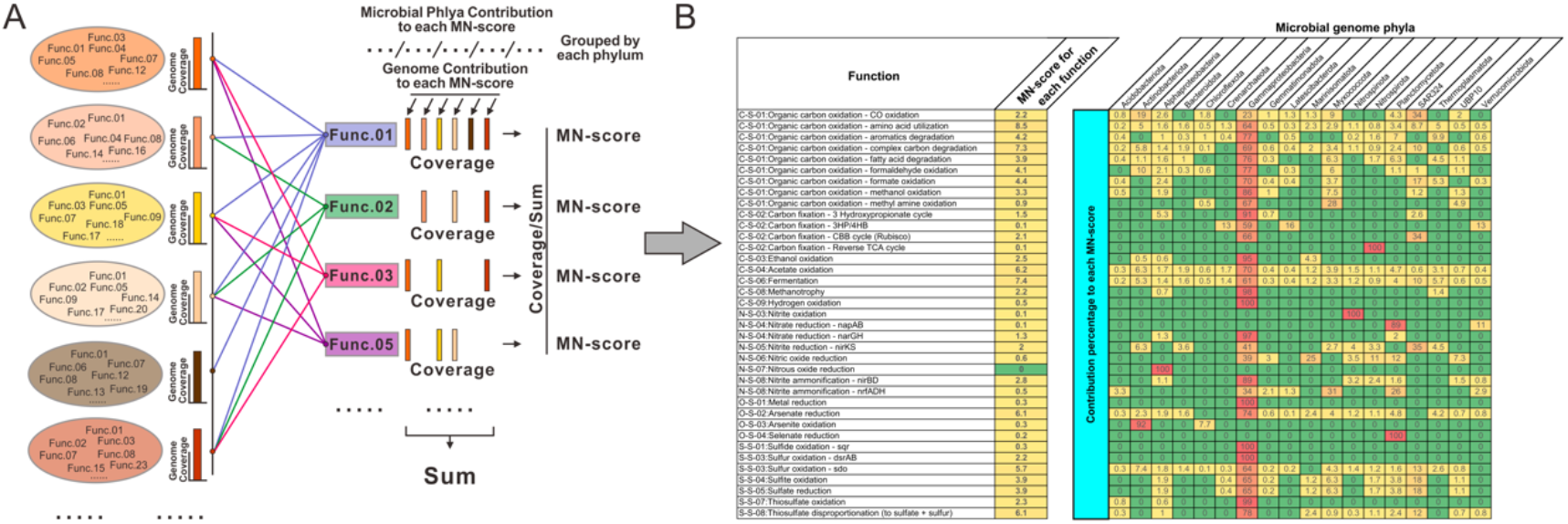
The calculation and result table of MN-score. **(A)** The calculation method for the MN-score within a community based on a given metagenomic dataset. Each circle stands for a genome within the community, and the adjacent bar stands for its genome coverage within the community. The coverage values of encoded genes for individual functions were summed up as the denominator, and the coverage value of encoded genes for each function was used as the numerator, and the MN-score was calculated accordingly for each function. **(B)** The resulted table of MN-score for the deep-sea hydrothermal vent metagenomic dataset. MN-score for each function was given in a separated column, and the rest part of the table indicates the contribution percentage to each MN-score of the genomes within the community as grouped by each phylum.

### Visualizing energy flow potential of metabolic reactions driven by microbial groups

To understand the contributions of microbial groups towards energy flow potential associated with specific metabolic and biogeochemical transformations, we developed an approach to visualize energy flow potential in communities at multiple scales including specific taxonomic groups, associated with a specific metabolic transformation, and entire biogeochemical cycles such as for carbon, nitrogen, or sulfur. Our approach involves the use of Sankey diagrams (also called ‘*Alluvial*’ plots) to represent the fractions of metabolic functions that are contributed by various microbial groups in a given community (Figure 6). This is referred to as an ‘energy flow potential’ diagram and allows visualization of metabolic reactions as the link between microbial contributors clustered as taxonomic groups and biogeochemical cycles at a community level (Figure 6 and Additional file 10: Dataset S3). The function fraction was calculated by accumulating the genome coverage values of genomes from a specific microbial group that possesses a given functional trait. The width of curved lines from a specific microbial group to a given functional trait indicates their corresponding proportional contribution to a specific metabolism (Figure 6). Alternatively, the genomic/metagenomic datasets which are used in constructing the above two diagrams: metabolic network diagram (Figure 4) and metabolic energy flow potential diagram (Figure 6), can be replaced by transcriptomic/metatranscriptomic datasets, and correspondingly, the gene coverage values will be replaced by gene expression values, and therefore, they will be representing the transcriptional activity patterns of metabolic network and metabolic energy flow potential at the community level (Additional file 2, 3, 4, and 5: Figure S2, S3, S4, and S5).

**Figure 6.**
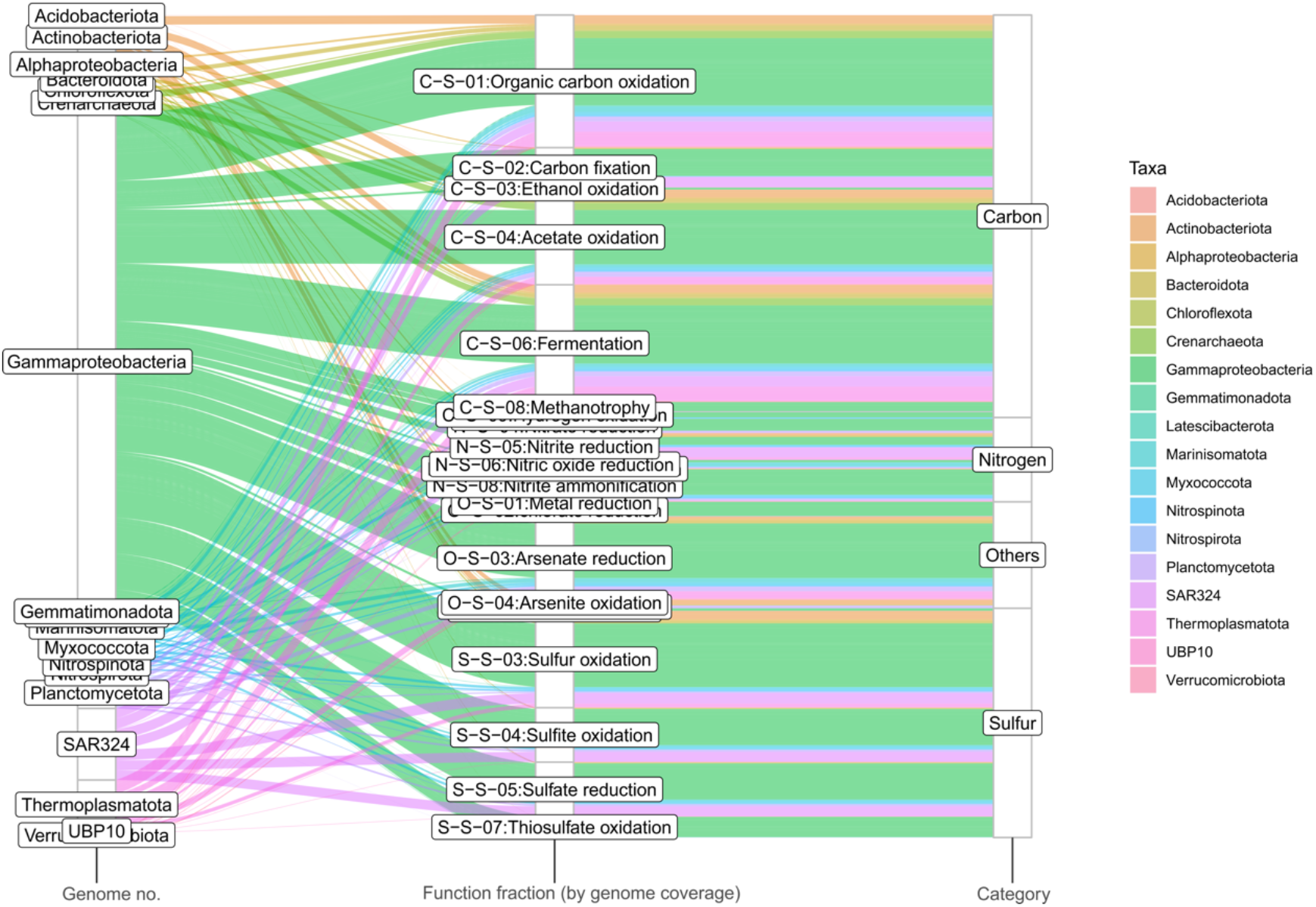
Metabolic energy flow potential diagram representing the contributions of microbial genomes to individual metabolic and biogeochemical processes, and at the scale of entire elemental cycles. Microbial genomes are represented at the phylum-level resolution. The three columns from left to right represent taxonomic groups scaled by the number of genomes, the contribution to each metabolic function by microbial groups calculated based on genome coverage, and the function category/biogeochemical cycle. The colors were assigned based on the taxonomy of the microbial groups. The deep-sea hydrothermal vent dataset was used for these analyses.

The microbial community dataset of 98 MAGs from a deep-sea hydrothermal vent was used as a test to demonstrate this workflow. After running the bioinformatic analyses described above, resulting tables and diagrams were compiled and visualized accordingly (Additional file 10: Dataset S3). Results for metabolic networks and MN-scores of the deep-sea hydrothermal vent environment indicate that the microbial community depends on mixotrophy and sulfur oxidation for energy conservation and involves in arsenate reduction potentially responsible for detoxification/arsenate resistance [57]. MN-scores indicate that amino acid utilization, complex carbon degradation, acetate oxidation, and fermentation are the major heterotrophic metabolisms for this environment; CO_2_-fixation and sulfur oxidation also occupy a considerable functional fraction, which indicates heterotrophy and autotrophy both contribute to energy conservation (Figure 5). Gammaproteobacteria are the most numerically abundant group in the community and they occupy significant functional fractions amongst both heterotrophic and autotrophic metabolisms (MN-score contribution ranging from 59-100%) (Figure 5, 6), which is consistent with previous findings in the Guaymas Basin hydrothermal environment. Meanwhile, MN-scores also explicitly reflect the involvement of other minor electron donors in energy conservation which are mainly contributed by Gammaproteobacteria, such as hydrogen and methane (Figure 5). This is also consistent with previous findings [43, 58] and indicates the accuracy and sensitivity of MN-scores to reflect metabolic potentials.

### METABOLIC is scalable, fast, and accurate

To test METABOLIC’s performance, we applied the software to analyze the metagenomic dataset which includes 98 MAGs from a deep-sea hydrothermal vent, and two sets of metagenomic reads (that are subsets of original reads with 10 million reads for each pair comprising ∼10% of the total reads). The total run time was ∼3 hours using 40 CPU threads in a Linux version 4.15.0-48-generic server (Ubuntu v5.4.0). The most compute-demanding part is hmmsearch, which took ∼45 mins. When tested on another dataset comprising ∼3600 microbial genomes (data not shown), METABOLIC could complete hmmsearch in ∼5 hours by using 40 CPU threads.

In order to test the accuracy of the results predicted by METABOLIC, we picked 15 bacterial and archaeal genomes from Chloroflexi, Thaumarchaeota, and Crenarchaeota which are reported to have 3 Hydroxypropionate cycle (3HP) and/or 3-hydroxypropionate/4-hydroxybutyrate cycle (3HP/4HB) for carbon fixation. METABOLIC predicted results in line with annotations from the KEGG genome database which can be visualized in KEGG Mapper (Table 1). Our predictions are also in accord with biochemical evidence of the existence of corresponding carbon fixation pathways in each microbial group: 1) 3 out of 5 *Chloroflexi* genomes are predicted by both METABOLIC and KEGG to possess the 3HP pathway and none of all these *Chloroflexi* genomes are predicted to possess the 3HP/4HB pathway. This is consistent with current reports based on biochemical and molecular experiments that only organisms from the phylum *Chloroflexi* are known to possess the 3HP pathway [59] (Table 1). 2) All 5 *Thaumarchaeota* genomes and 2 out of 5 *Crenarchaeota* genomes are predicted by both METABOLIC and KEGG to possess the 3HP/4HB pathway and none of these *Thaumarchaeota* and *Crenarchaeota* genomes are predicted to possess the 3HP pathway. This is consistent with current reports that only the 3HP/4HB pathway could be detected in *Crenarchaeota* and *Thaumarchaeota* [60, 61] (Table 1). We have also applied METABOLIC on a large well-studied dataset comprising 2545 metagenome-assembled genomes from terrestrial subsurface sediments and groundwater [2]. The annotation results of METABOLIC are consistent with previously described reports (Additional file 6, 10: Figure S6, Dataset S3). These results suggest that METABOLIC can provide accurate annotations and genomic profiles and perform well as a functional predictor for microbial genomes and communities.

**Table 1.**
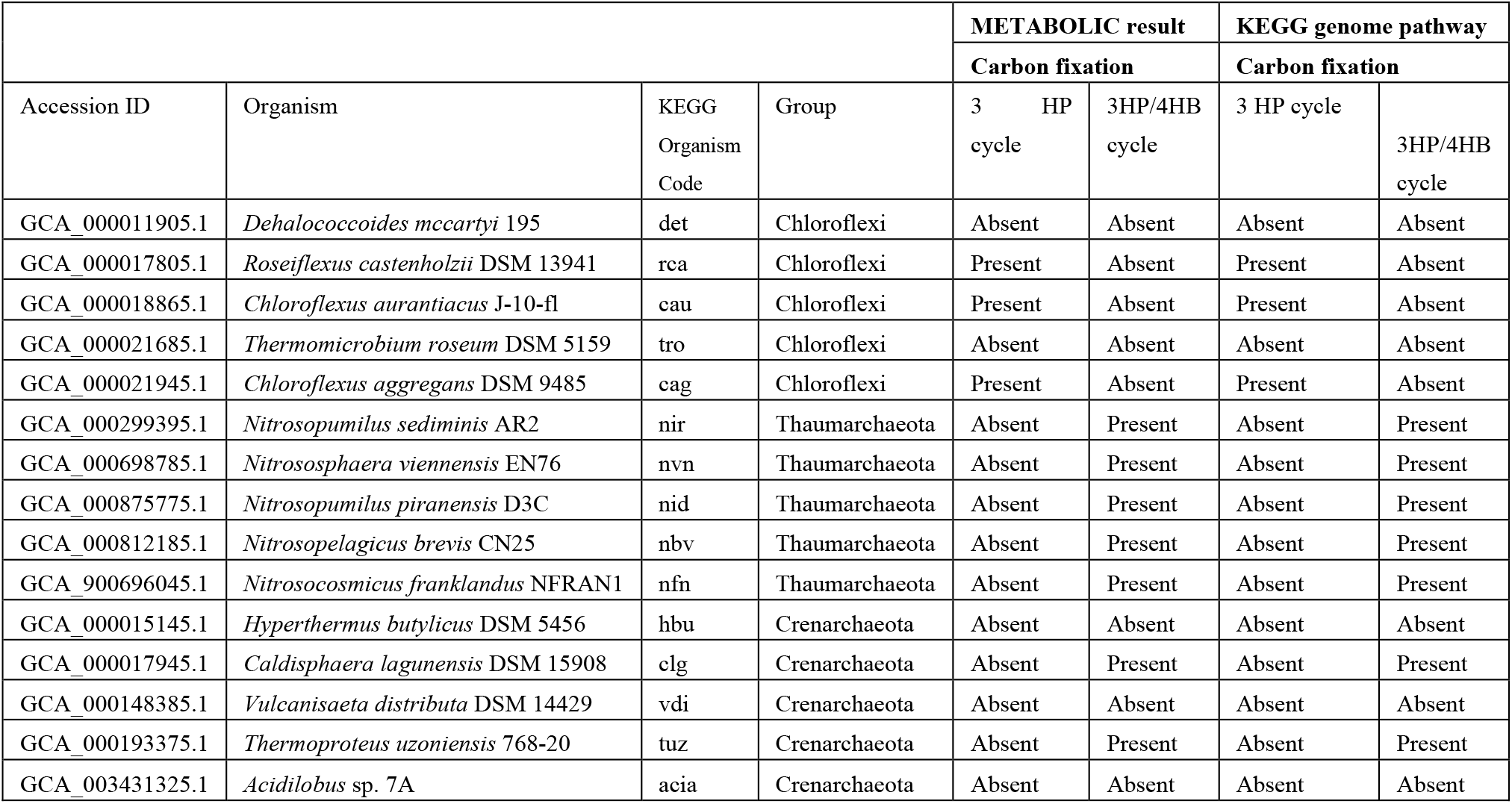
The carbon fixation metabolic traits of 15 tested bacterial and archaeal genomes predicted by both METABOLIC and KEGG genome database

|  |  |  |  | METABOLIC result |  | KEGG genome pathway |  |
| --- | --- | --- | --- | --- | --- | --- | --- |
|  |  |  |  | Carbon fixation |  | Carbon fixation |  |
| Accession ID | Organism | KEGG Organism Code | Group | 3 HP cycle | 3HP/4HB cycle | 3 HP cycle | 3HP/4HB cycle |
| GCA_000011905.1 | *Dehalococcoides mccartyi* 195 | det | Chloroflexi | Absent | Absent | Absent | Absent |
| GCA_000017805.1 | *Roseiflexus castenholzii* DSM 13941 | rca | Chloroflexi | Present | Absent | Present | Absent |
| GCA_000018865.1 | *Chloroflexus aurantiacus* J-10-fl | cau | Chloroflexi | Present | Absent | Present | Absent |
| GCA_000021685.1 | *Thermomicrobium roseum* DSM 5159 | tro | Chloroflexi | Absent | Absent | Absent | Absent |
| GCA_000021945.1 | *Chloroflexus aggregans* DSM 9485 | cag | Chloroflexi | Present | Absent | Present | Absent |
| GCA_000299395.1 | *Nitrosopumilus sediminis* AR2 | nir | Thaumarchaeota | Absent | Present | Absent | Present |
| GCA_000698785.1 | *Nitrososphaera viennensis* EN76 | nvn | Thaumarchaeota | Absent | Present | Absent | Present |
| GCA_000875775.1 | *Nitrosopumilus piranensis* D3C | nid | Thaumarchaeota | Absent | Present | Absent | Present |
| GCA_000812185.1 | *Nitrosopelagicus brevis* CN25 | nbv | Thaumarchaeota | Absent | Present | Absent | Present |
| GCA_900696045.1 | *Nitrosocosmicus franklandus* NFRAN1 | nfn | Thaumarchaeota | Absent | Present | Absent | Present |
| GCA_000015145.1 | *Hyperthermus butylicus* DSM 5456 | hbu | Crenarchaeota | Absent | Absent | Absent | Absent |
| GCA_000017945.1 | *Caldisphaera lagunensis* DSM 15908 | clg | Crenarchaeota | Absent | Present | Absent | Present |
| GCA_000148385.1 | *Vulcanisaeta distributa* DSM 14429 | vdi | Crenarchaeota | Absent | Absent | Absent | Absent |
| GCA_000193375.1 | *Thermoproteus uzoniensis* 768-20 | tuz | Crenarchaeota | Absent | Present | Absent | Present |
| GCA_003431325.1 | *Acidilobus* sp. 7A | acia | Crenarchaeota | Absent | Absent | Absent | Absent |

### METABOLIC provides robust performance and consistent metabolic analyses

Currently, several software packages and online servers are available for genome annotation and metabolic profiling. However, METABOLIC is unique in its ability to integrate multi-omic information towards elucidating metabolic connections, energy flow, and contribution of microorganisms to biogeochemical cycles. We compared the performance of METABOLIC (Figure 7A) to other software including GhostKOALA [62], BlastKOALA [62], KAAS [63], and RAST/SEED [22].

**Figure 7.**
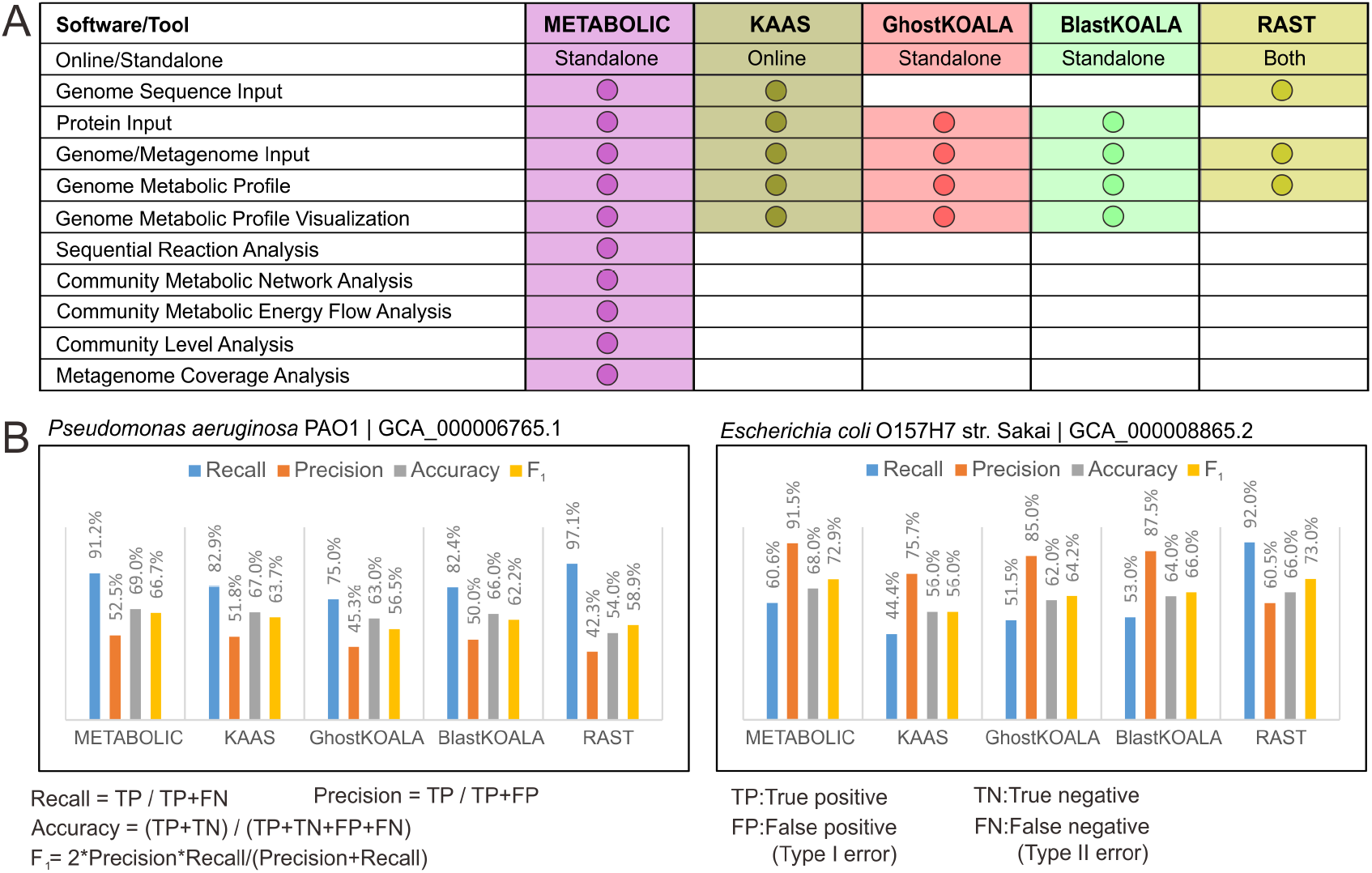
Comparison of METABOLIC with other software packages and online servers. **(A)** Comparison of the workflows and services, **(B)** Comparison of performance of protein prediction for two representative genomes, *Pseudomonas aeruginosa* PAO1, and *Escherichia coli* O157H7 str. sakai.

To compare the prediction performance (Figure 7B), we used two representative bacterial genomes as the test datasets. We randomly picked 100 protein sequences from individual genomes and submitted them to annotation by these five software/online servers. Predicted protein annotations by individual software and online servers were compared to their original annotations that were provided by the NCBI database (Additional file 11, 12: Dataset S4, S5). According to statistical methods of binary classification [64], the following parameters were used to make the comparison: 1) recall (also referred to as the sensitivity) as the true positive rate, 2) precision (also referred to as the positive predictive value) which indicates the reproducibility and repeatability of a measurement system, 3) accuracy which indicates the closeness of measurements to their true values, and 4) F1 value which is the harmonic mean of precision and recall, and reflects both these two parameters. Among the tested software/servers, the performance parameters of METABOLIC consistently placed it in the top 2 programs for recall and F1 and as the best for precision and accuracy. These results demonstrate that METABOLIC (Figure 7B) provides robust performance and consistent metabolic prediction for genomes that offer wide applicability of use for the downstream visualization and community-level analysis.

### Metabolic and biogeochemical comparisons at the community scale in diverse environments

To demonstrate the application and performance of METABOLIC in different samples, we tested eight distinct environments (marine subsurface, terrestrial subsurface, deep-sea hydrothermal vent, freshwater lake, gut microbiome from patients with colorectal cancer, gut microbiome from healthy control, meadow soil, wastewater treatment facility). Overall, we found METABOLIC to perform well across all the environments to profile microbial genomes with functional traits and biogeochemical cycles (Additional file 10: Dataset S3). Within these tested environments, we also performed community-scale metabolic comparisons based on the MN-score (Figure 8). MN-score fraction at the community scale reflects the overall metabolic profile distribution. Specifically, we compared samples from terrestrial and marine subsurface and samples from hydrothermal vent and freshwater lake. We observed that terrestrial subsurface contains more abundant metabolic functions related to nitrogen cycling compared to the marine subsurface (Figure 8A), consistent with the previous characterization of these two environments [2, 65]. Deep-sea hydrothermal vent samples had a considerably high concentration of methane and hydrogen [43] as compared to Lake Tanganyika (freshwater lake); the deep-sea hydrothermal vent microbial community has more abundant metabolic functions associated with methanotrophy and hydrogen oxidation (Figure 8B). To focus on a specific biogeochemical cycle, we applied METABOLIC to compare sulfur related metabolisms at the community scale for these two environment pairs (Additional file 7: Figure S7). Terrestrial subsurface contains genomes covering more sulfur cycling steps compared to marine subsurface (7 steps vs 3 steps) (Additional file 7: Figure S7A). Freshwater lake contains genomes involving almost all the sulfur cycling steps except for sulfur reduction, while deep-sea hydrothermal vent contains less sulfur cycling steps (8 steps vs 6 steps) (Additional file 7: Figure S7B). Nevertheless, deep-sea hydrothermal vent has a higher fraction of genomes (59/98) and a higher relative abundance (73%) of these genomes involving sulfur oxidation compared to the freshwater lake (Additional file 7: Figure S7B). This indicates that the deep-sea hydrothermal vent microbial community has a more biased sulfur metabolism towards sulfur oxidation, which is consistent with previous metabolic characterization on the dependency of elemental sulfur in this environment [43, 66-68]. Collectively, by characterizing community-scale metabolism, METABOLIC can facilitate the comparison of overall functional profiles as well as functional profiles for a particular elemental cycle.

**Figure 8.**
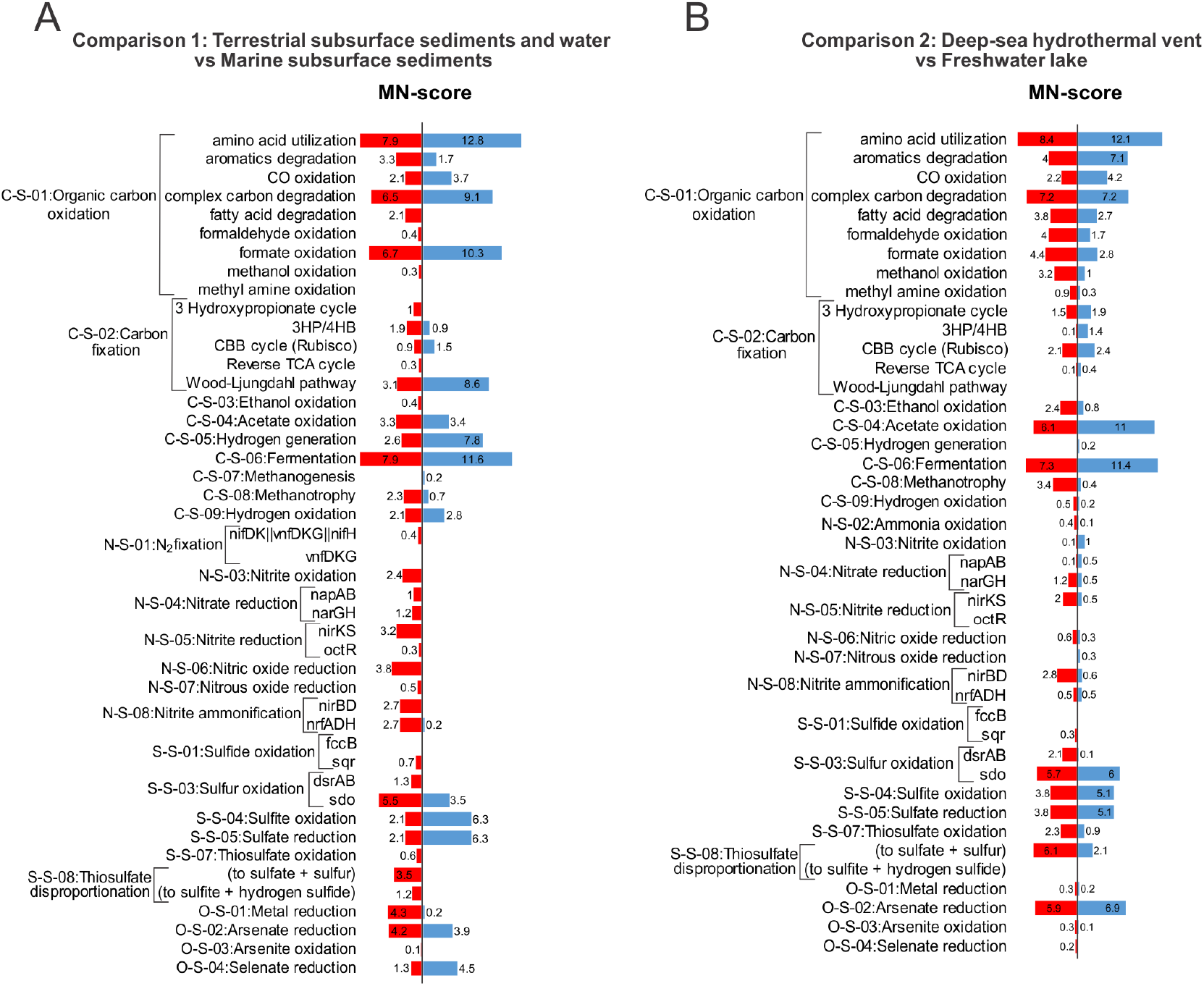
Community metabolism comparison based on MN-scores. **(A)** Comparison between marine subsurface and terrestrial subsurface. **(B)** Comparison between freshwater lake and deep-sea hydrothermal vent. MN-scores were calculated as gene coverage fractions for individual metabolic functions. Functions with MN-scores in both environments as zero were removed from each panel, e.g., N-S-02:Ammonia oxidation, N-S-09:Anammox, S-S-02:Sulfur reduction, and S-S-06:Sulfite reduction in Panel (A), and C-S-07:Methanogenesis, N-S-01:N2 fixation, N-S-09:Anammox, S-S-02:Sulfur reduction, and S-S-06:Sulfite reduction in Panel (B). Details for MN-score and each microbial group contribution refer to Supplementary Dataset S3.

### METABOLIC enables accurate reconstruction of cell metabolism

To demonstrate applications of reconstructing and depicting cell metabolism based on METABOLIC results, two microbial genomes were used as an example (Figure 9). As illustrated in Figure 9A, Hadesarchaea archaeon 1244-C3-H4-B1 has no TCA cycling gene components, which is consistent with previous findings in archaea within this class [69]. Gluconeogenesis/glycolysis pathways are also lacking in the genome; since gluconeogenesis is the central carbon metabolism responsible for generating sugar monomers which will be further biosynthesized to polysaccharides as important cell structural components [70], the lack of this pathway could be due to genome incompleteness. As an enigmatic archaeal class newly discovered in the recent decade, Hadesarchaea have distinctive metabolisms that separate them from conventional euryarchaeotal groups. They almost lost all TCA cycle gene components for the production of acetyl-CoA; while they could metabolize amino acids in a heterotrophic lifestyle [69]. It is posited that the Hadesarchaea genome has been subjected to streamline processing possibly due to nutrient limitations in their surrounding environment [69]. Due to their metabolic novelty and limited available genomes in the current time, there are still uncertainties on unknown/hypothetical genes and pathways and unclassified metabolic potential across the whole class. The previous metabolic characterization on four Hadesarchaea genomes indicates Hadesarchaea members could anaerobically oxidize CO and H2 was produced as the side product [69]. In the Hadesarchaea archaeon 1244-C3-H4-B1 genome, METABOLIC results indicate the loss of all anaerobic carbon-monoxide dehydrogenase gene components, which suggests the distinctive metabolism of this Hadesarchaea archaeon from others and highlights the accuracy of METABOLIC in reflecting functional details.

**Figure 9.**
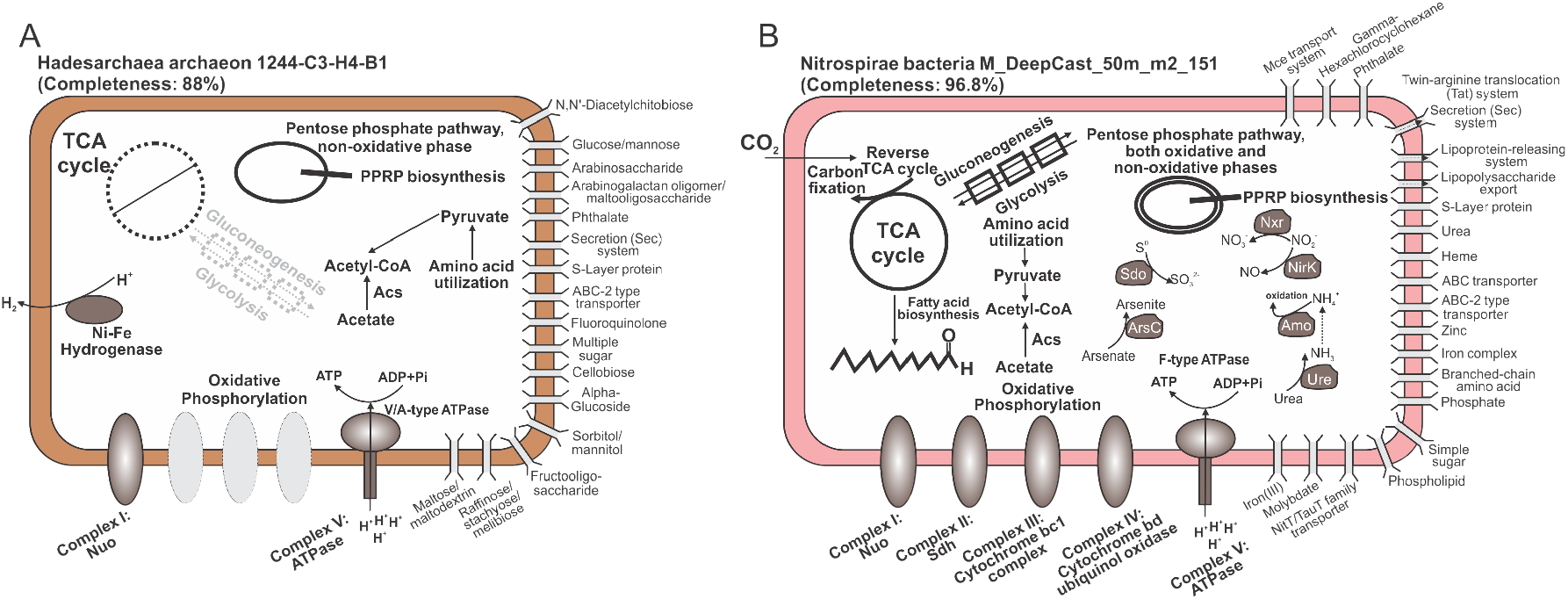
Cell metabolism diagrams of two microbial genomes. **(A)** cell metabolism diagram of Hadesarchaea archaeon 1244-C3-H4-B1 **(B)** cell metabolism diagram of Nitrospirae bacteria M_DeepCast_50m_m2_151. The absent functional pathways/complexes were labeled with dash lines.

We also reconstructed the metabolism for Nitrospirae bacteria M_DeepCast_50m_m2_151, a member of the *Nitrospirae* phylum reconstructed from Lake Tanganyika [46] (Figure 9B), it contains the full pathway for the TCA cycle and gluconeogenesis/glycolysis. Furthermore, it also has the full set of oxidative phosphorylation complexes for energy conservation and functional genes for nitrite oxidation to nitrate. Other nitrogen cycling metabolisms identified in this genome include ammonium oxidation, urea utilization, and nitrite reduction to nitric oxide. The Reverse TCA cycle pathway was identified for carbon fixation. The metabolic profiling result is in accord with the fact that Nitrospirae is a well-known nitrifying bacterial class capable of nitrite oxidation and living an autotrophic lifestyle [70]. Additionally, their more abundant distribution in nature compared to other nitrite-oxidizing bacteria such as *Nitrobacter* indicates a significant contribution to nitrogen cycling in the environment [70]. This highlights the ability of METABOLIC in reflecting functional details of more common and prevalent microorganisms compared to the Hadesarchaea archaeon. Notably as discovered from METABOLIC analyses, this bacterial genome also contains a wide range of transporter enzymes on the cell membrane, including mineral and organic ion transporters, sugar and lipid transporters, phosphate and amino acid transporters, heme and urea transporters, lipopolysaccharide and lipoprotein releasing system, bacterial secretion system, etc., which indicates its metabolic versatility and potential interactive activities with other organisms and the ambient environment. Collectively, METABOLIC result of functional profiling provides an intuitively-represented summary of a single microbial genome which enables depicting cell metabolism for better visualization of the functional capacity.

### METABOLIC accurately represents metabolism in the human microbiome

In addition to resolving microbial metabolism and biogeochemistry in environmental microbiomes, METABOLIC also accurately identifies metabolic traits associated with human microbiomes. The human microbiome contributes to normal human development, human physiology, and disease pathology. Study of human microbiomes are an advancing field and has been accelerated by the NIH’s implementation of Human Microbiome Project [71]. While healthy and disease state human microbiome samples continue to be collected and sequenced at a rapid pace, the implications of microbial metabolism on human health largely remain a black box, much like microbial contributions to biogeochemical cycling. We demonstrate the utility of METABOLIC in highlighting metabolism in human microbiomes using publicly available samples from a study of human microbiome in colorectal cancer using stool samples collected from patients with colorectal cancer and healthy individuals. From the study, we selected one colorectal cancer (CRC) and an age and sex matched control (healthy human) gut metagenomes from stool samples to conduct the comparison (Figure 10). The heatmap indicates the human microbiome functional profiles of both samples based on the marker gene presence/absence patterns (Figure 10). As an example of METABOLIC’s application, we demonstrate that there were 28 makers with variations > 10% in terms of the marker-containing genome numbers between these two states (Figure 10). These 28 markers involved all the ten metabolic categories except for lipid metabolism and translation, suggesting the broad functional differences between these two states. In addition to analyzing the human microbiome specific-functional markers, METABOLIC can be used as described in previous sections on human microbiome samples to visualize elemental nutrient cycling and analyze metabolic nutrient interaction. METABOLIC results provide a comprehensive functional profile that could be to represent human-microbial interactions; overall it enables systematic characterization of the composition, structure, function, and dynamics of microbial metabolisms in the human microbiome and facilitates omics-based studies of microbial community on human health [50].

**Figure 10.**
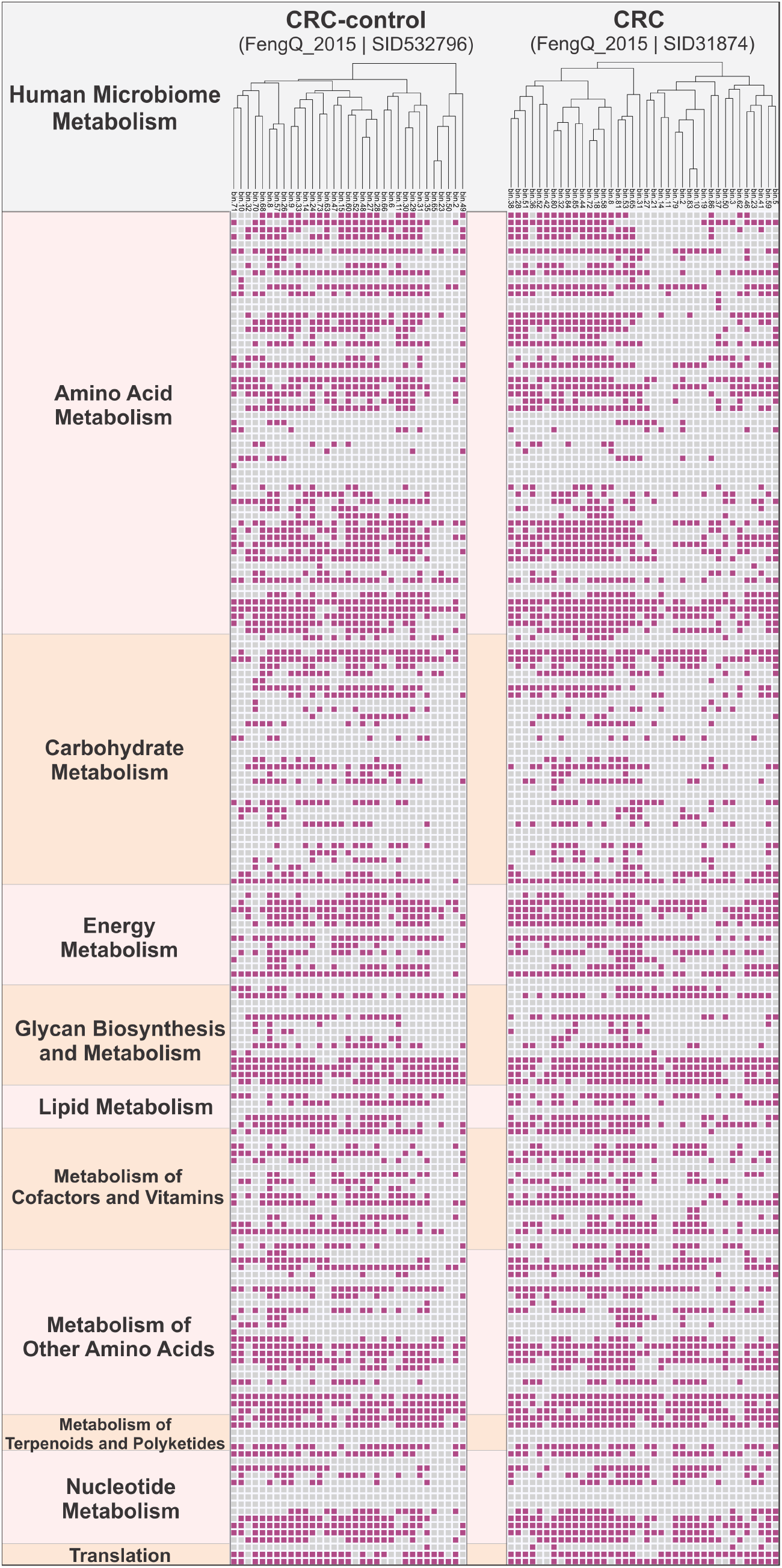
Presence/Absence map of human microbiome metabolisms of a colorectal cancer patient (CRC) and a healthy control gut samples. The heatmap has summarized 189 horizontal entries (189 lines) from 139 key functional gene families that covered 10 function categories. Detailed KEGG KO identifier IDs and protein information for each function category were described in Supplementary Dataset S2.

## Conclusions

In the recent decade, the rapidly growing number of sequenced microbial genomes, including pure isolates, metagenome-assembled genomes, and single-cell genomes, have significantly contributed to the growth of microbial genome databases, which has made large-scale microbial genome analyses more tractable. Metabolic functional profile of microbial genomes at the scale of individual organisms and communities is essential for microbial ecologists and biogeochemists to have a comprehensive understanding of ecosystem processes and biogeochemistry, and as a conduit for enabling trait-based models of biogeochemistry. We have developed METABOLIC as a metabolic functional profiler that goes above and beyond current frameworks of genome/protein annotation platforms in providing protein annotations and metabolic pathway analyses that are used for inferring contribution of microorganisms, metabolism, interactions, activity, and biogeochemistry at the community-scale. METABOLIC is the first software to enable community-scale visualization of microbial metabolic handoffs, interactions, and contributions to biogeochemical cycles. We anticipate that METABOLIC will enable easier interpretation of microbial metabolism and biogeochemistry from metagenomes and genomes and enable microbiome research in diverse fields. Finally, METABOLIC will facilitate standardization and integration of genome-informed metabolism into metabolic and biogeochemical models.

## Additional files

**Additional file 1: Figure S1**. METABOLIC result tables

**Additional file 2: Figure S2**. Metabolic network diagram based on the transcriptomic dataset from a hydrothermal vent sample

**Additional file 3: Figure S3**. Metabolic network diagram based on the transcriptomic dataset from hydrothermal background sample

**Additional file 4: Figure S4**. Microbial metabolic energy flow potential diagram based on the transcriptomic dataset from hydrothermal vent sample

**Additional file 5: Figure S5**. Microbial metabolic energy flow potential diagram based on the transcriptomic dataset from hydrothermal background sample

**Additional file 6: Figure S6**. Metabolic profile diagram of terrestrial subsurface microbial community

**Additional file 7: Figure S7**. Comparison of sulfur related metabolism at the community scale level

**Additional file 8: Dataset S1**. The motif sequences and motif pairs

**Additional file 9: Dataset S2**. Summary table of Human Microbiome Marker genes

**Additional file 10: Dataset S3**. METABOLIC result of eight different environments

**Additional file 11: Dataset S4**. The comparison of the protein prediction performance among five software packages/online servers on the genome of *Escherichia coli* O157H7 str. Sakai

**Additional file 12: Dataset S5**. The comparison of the protein prediction performance among five software packages/online servers on the genome of *Pseudomonas aeruginosa* PAO1

## Acknowledgments

We thank the comments and suggestions from the users of METABOLIC, which helped to improve and expand the functions of this software.

## Authors’ contributions

ZZ and KA conceptualized and designed the study. ZZ and PQT wrote the Perl and R scripts. ZZ ran the test data and improved the software. YL provided a part of the databases. PQT, AMB, KK, ESC, and UK provided ideas and comments, helped to set up the GitHub page, and contributed to improving the overall performance of the software. ZZ and KA wrote the manuscript, and all authors contributed and approved the final edition of the manuscript.

## Corresponding authors

Correspondence to Karthik Anantharaman.

## Ethics declarations

### Ethics approval and consent to participate

Not applicable.

## Consent for publication

Not applicable.

