## Supplementary Figure S1 for "METABOLIC: High-throughput profiling of microbial genomes for functional traits, biogeochemistry, and community-scale metabolic networks"

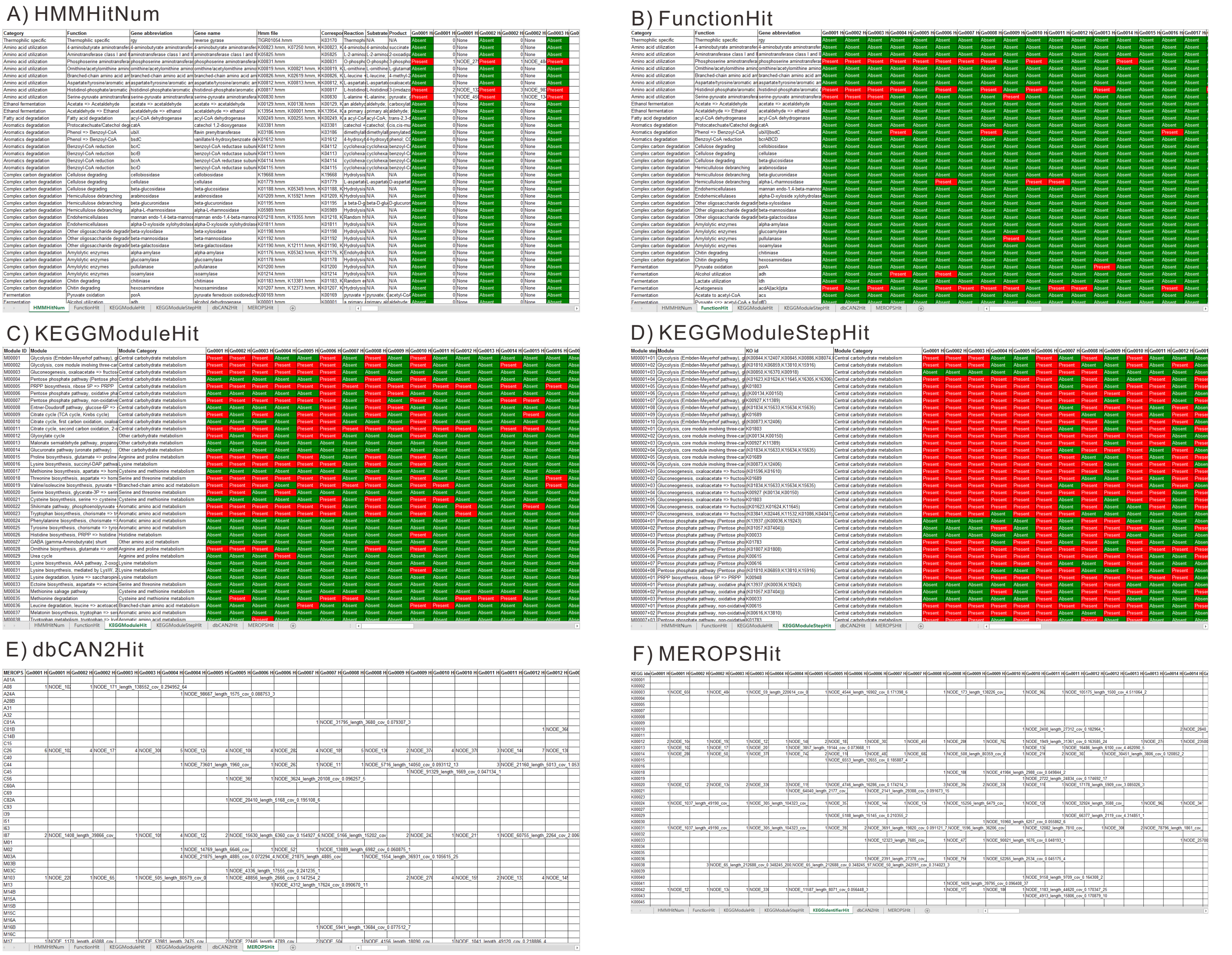


**Supplementary Figure S1. METABOLIC result table report.** A) “HMMHitNum” worksheet represents the function detail information and reflects the presence/absence of an HMM profile, HMM hit protein numbers and IDs. B) “FunctionHit” worksheet reflects the presence/absence of a functional trait. C) “KEGGModuleHit” worksheet reflects the presence/absence of a KEGG module by given a module cutoff value. D) “KEGGModuleStepHit” worksheet reflects the presence/absence of a KEGG module step. E) “dbCAN2Hit” worksheet reflects the hit numbers and names of CAZymes. F) “MEROPSHit” worksheet reflects the hit numbers and names of peptidases and inhibitors.
