## Supplementary Figure S2 for "METABOLIC: High-throughput profiling of microbial genomes for functional traits, biogeochemistry, and community-scale metabolic networks"

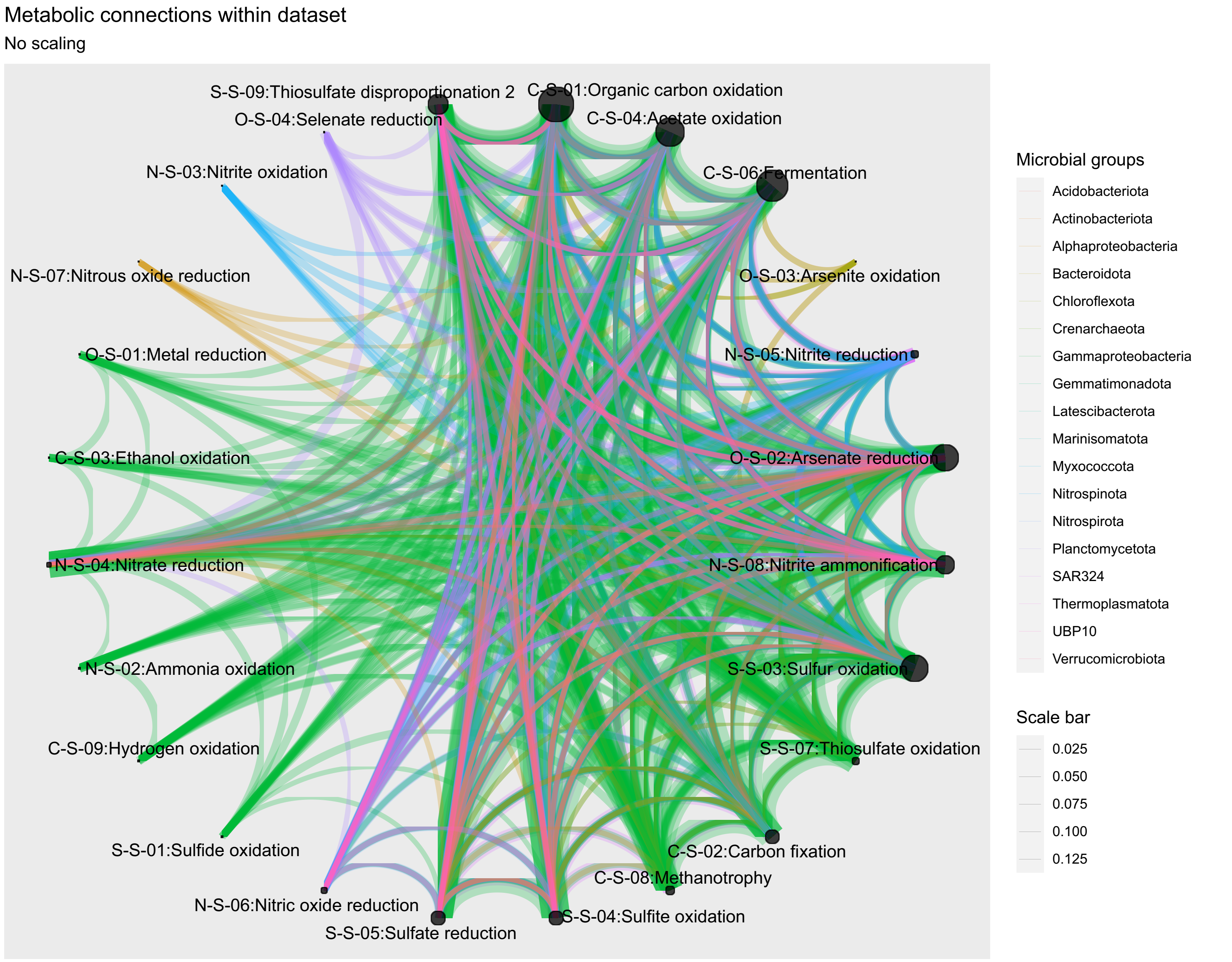


**Supplementary Figure S2. Metabolic network diagram based on the transcriptomic dataset from hydrothermal vent sample.** The nodes represent biogeochemical cycling steps, and edges connecting two given nodes represent the metabolic connections between them. The thickness of the edge was depicted according to the average of the gene expression values of the connected two biogeochemical cycling steps, which were calculated by the transcriptomic dataset from Guaymas Basin hydrothermal vent sample. The color of the edge was assigned based on the taxonomic group of the represented genome.
