## Supplementary Figure S5 for "METABOLIC: High-throughput profiling of microbial genomes for functional traits, biogeochemistry, and community-scale metabolic networks"

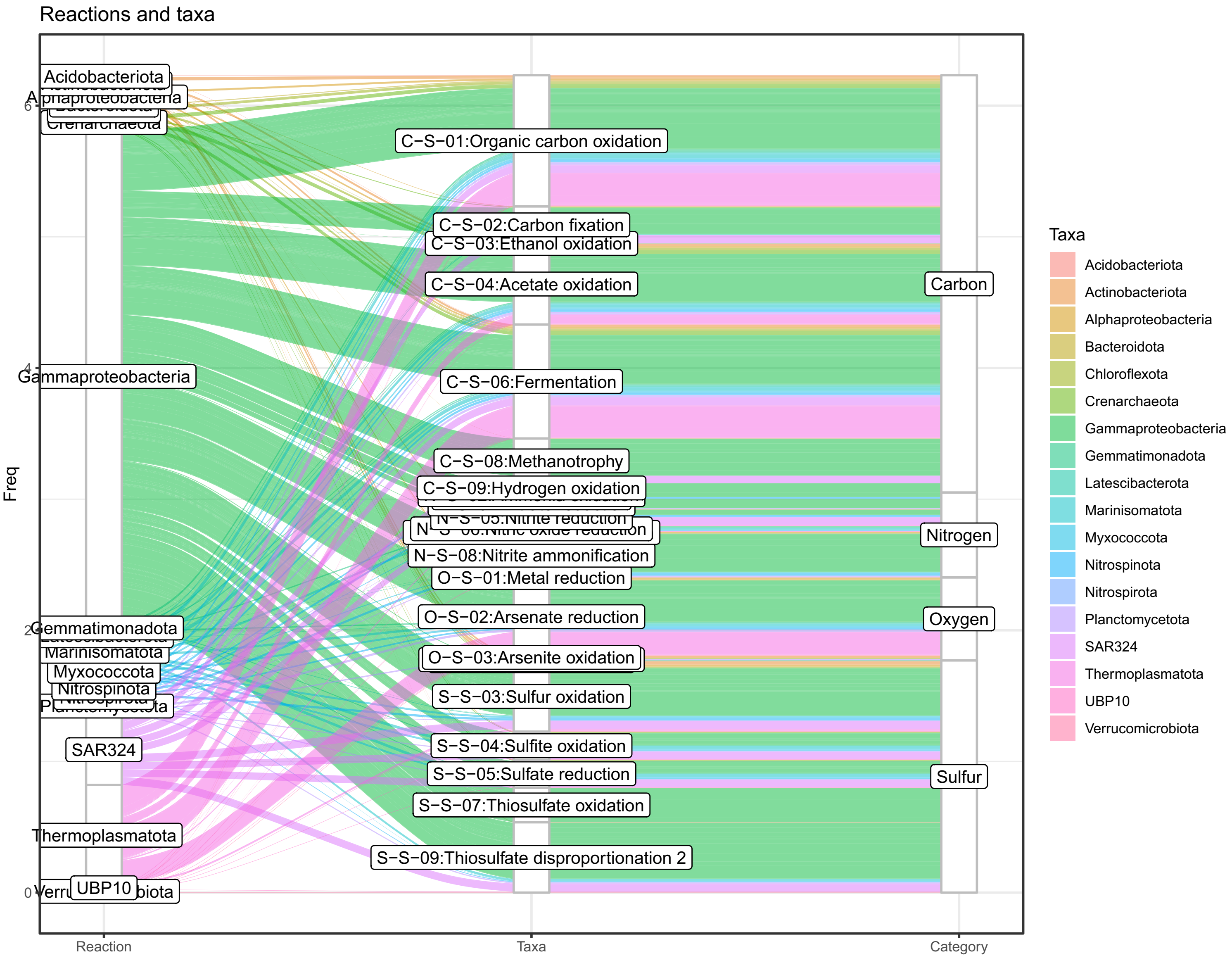


**Supplementary Figure S5. Microbial metabolic energy flow diagram based on the transcriptomic dataset from hydrothermal background sample.** The three columns of the network diagram represent taxonomic groups and genome numbers, the expression fraction of each microbial group calculated by transcriptome coverage, and the function category.
