## Supplementary Figure S6 for "METABOLIC: High-throughput profiling of microbial genomes for functional traits, biogeochemistry, and community-scale metabolic networks"

| | Genome Group ID | G01 | G02 | G03 | G04 | G05 | G06 | G07 | G08 | G09 | G10 | G11 | G12 | G13 | G14 | G15 | G16 | G17 | G18 | G19 | G20 | G21 | G22 | G23 | G24 | G25 | G26 | G27 | G28 | G29 | G30 | G31 | G32 | G33 | G34 | G35 | G36 | G37 | G38 | G39 | G40 |
|---|---|---|---|---|---|---|---|---|---|---|---|---|---|---|---|---|---|---|---|---|---|---|---|---|---|---|---|---|---|---|---|---|---|---|---|---|---|---|---|---|---|
| | Genome Number | 12 | 3 | 16 | 3 | 36 | 7 | 1 | 6 | 6 | 4 | 8 | 4 | 1 | 19 | 1 | 1 | 5 | 5 | 61 | 1 | 1 | 1 | 3 | 7 | 2 | 2 | 3 | 8 | 1 | 9 | 6 | 7 | 2 | 1 | 9 | 2 | 5 | 1 | 11 | 2 |
| | Thermophilic speci c | | | | | | | | | | | | | | | | | | | | | | | | | | | | | | | | | | | 1 | | | | | |
| C metabolism | Amino acid utilization | 11 | 3 | 15 | 2 | 36 | 4 | 1 | 6 | 5 | 3 | 8 | 4 | 1 | 19 | 1 | 1 | 5 | 5 | 61 | 1 | 1 | 1 | 3 | 6 | 2 | 2 | 3 | 8 | | 9 | | 7 | 2 | 1 | 9 | 2 | 4 | 1 | 10 | 1 |
| C metabolism | Aromatics degradation | 9 | 1 | 8 | 2 | 20 | 1 | | 6 | 2 | 3 | 7 | 4 | | 4 | | | 1 | 5 | 46 | 1 | | 1 | 2 | | 2 | 1 | 3 | 6 | | 4 | | 2 | 2 | 1 | 1 | 1 | 3 | 1 | 2 | 1 |
| C metabolism | Complex carbon degradation | 7 | 1 | 10 | 2 | 29 | 3 | 1 | | 1 | | 3 | 1 | | 4 | 1 | 1 | 3 | 3 | 27 | 1 | 1 | 1 | 3 | | 2 | | | 2 | 1 | | 1 | 4 | | | 3 | | | 1 | 10 | 1 |
| C metabolism | Fermentation | 11 | 3 | 7 | 3 | 35 | 7 | 1 | 6 | 6 | 2 | 8 | 4 | 1 | 19 | 1 | 1 | 5 | 5 | 44 | 1 | 1 | 1 | 3 | 7 | 2 | 2 | 3 | 8 | 1 | 7 | 3 | 6 | 2 | 1 | 9 | 2 | 1 | 1 | 10 | 2 |
| C metabolism | C1 metabolism\|Aerobic CO oxidation | 9 | 1 | 3 | 1 | 8 | 2 | 1 | | 4 | | | | | 9 | | | 1 | | 23 | | | 1 | | | 1 | | 1 | 1 | | 1 | | | | 1 | 1 | | | 1 | 1 | 1 |
| C metabolism | C1 metabolism\|Formaldehyde oxidation | | | 10 | | | | | | | | | | | | | | | | 32 | | | | | | | | | | | | | | | | | | | | | |
| C metabolism | $C_1$ metabolism\|Formate oxidation | 7 | 1 | 4 | 1 | 21 | 3 | 1 | | 4 | | 7 | 4 | | 1 | | | | | 32 | | 1 | 1 | 3 | 4 | 2 | 1 | 1 | 6 | | 2 | | 2 | | 1 | 5 | 2 | | 1 | 4 | 2 |
| C metabolism | $C_1$ metabolism\|Methanol oxidation | 4 | | 8 | | | | | 1 | | | | | | | | | | | 14 | | | | | | | | | | | | | | | | | | | | | |
| C metabolism | $C_1$ metabolism\|Methyl amine -> formaldehyde | | | 3 | | | | | | | | | | | | | | | | 1 | | | | | | | | | | | | | | | | | | | | | |
| C metabolism | Methane oxidation | | | 4 | | 2 | | | | | | | | | 3 | | | | 1 | 12 | | | | | 1 | | 1 | | | | 2 | 1 | 1 | | | | | | | 1 | 1 |
| C metabolism | Methane production | | | | | | | | | | | | | | | | | | | | | | | | | | | | | | | | | | | | | | | | |
| C metabolism | Carbon xation\|3 Hydroxypropionate cycle | 1 | | 4 | | 3 | | | | | | 3 | | | | | | | | 22 | | | | | | 1 | | | 2 | | | | | | | 2 | | | | | |
| C metabolism | Carbon xation\|3HP/4HB | 2 | 3 | 2 | 1 | 2 | | | | 1 | | 3 | | | | | | | | 1 | | | 1 | 1 | | 2 | | | | | 1 | | | | | 2 | | | | | 1 |
| C metabolism | Carbon xation\|CBB cycle - Rubisco | | | 10 | | | | | | | | | | | | | | | | 29 | | | | | | | | | | | | | | | | | | | | | |
| C metabolism | Carbon xation\|Reverse TCA cycle | | | | | | 1 | | 6 | | | | | | 7 | | | | | | | | | | | | | 2 | 3 | | | 1 | | | | | | | | | |
| C metabolism | Carbon xation\|Wood-Ljungdahl pathway | 1 | 2 | | 1 | 3 | 1 | | | 5 | 2 | 7 | 3 | | 8 | 1 | 1 | 1 | 3 | | | 1 | | 2 | 2 | | | | 5 | | 2 | | 5 | | | 6 | 1 | | | 7 | |
| N metabolism | Nitrogen cycling\|Ammonia oxidation | | | | | | | | | | | | | | | | | | | | | | | | | | | | | | | | | | | | | | | | |
| N metabolism | Nitrogen cycling\|Anammox | | | | | | | | | | | | | | | | | | | | | | | | | | | | | | | | 3 | | | | | | | | |
| N metabolism | Nitrogen cycling\|N2 fixation | | | | | | | | 4 | | 1 | 5 | 3 | | 2 | | | | | 8 | | | | | 1 | | 1 | | 5 | | | | | 2 | | 4 | | | 1 | 2 | |
| N metabolism | Nitrogen cycling\|Nitrate reduction | 5 | | 6 | 1 | 7 | 1 | | 5 | 2 | 1 | 4 | 2 | | 2 | | | | 3 | 33 | 1 | | | 2 | | 2 | 1 | | 2 | | 1 | | | 2 | 1 | | | 2 | | | |
| N metabolism | Nitrogen cycling\|Nitric oxide reduction | 8 | | 4 | 1 | 15 | 5 | | 5 | | | 3 | 1 | | 1 | | | | 1 | 23 | 1 | | 1 | 3 | | 2 | 1 | 1 | 4 | | 5 | | 2 | | 1 | 2 | | | 1 | | 2 |
| N metabolism | Nitrogen cycling\|Nitrite oxidation | 6 | | | 1 | 2 | | | | 1 | | | | | 3 | | | | 1 | 3 | | | 1 | | | 2 | 1 | 1 | | 1 | 6 | | 1 | | 1 | | 1 | 3 | 1 | 1 | 1 |
| N metabolism | Nitrogen cycling\|Nitrite reduction | 4 | 1 | 4 | | 4 | 4 | | 3 | 3 | | 1 | 2 | | | | | | 3 | 25 | | | | | | 1 | | 2 | 3 | 1 | 1 | 1 | 1 | | | | 1 | 3 | | | 1 |
| N metabolism | Nitrogen cycling\|Nitrite reduction to ammonia | 3 | | 8 | 2 | 21 | 4 | | | 2 | | 6 | 4 | | 8 | | | | 2 | 39 | 1 | 1 | 1 | 3 | | 1 | 2 | 1 | 8 | | 4 | | 2 | | 1 | 3 | 1 | 4 | | 3 | 1 |
| N metabolism | Nitrogen cycling\|Nitrous oxide reduction | 2 | | | 2 | 9 | | | 3 | 1 | | | | | 3 | | | | | 1 | | | | | 1 | | | | | | 1 | | | | | | | | | | |
| S metabolism | Sulfur cycling\|Sulfate reduction | 8 | 1 | 12 | 2 | 24 | 1 | | 5 | 2 | 3 | 8 | 2 | | 5 | | 1 | | 5 | 49 | | | 1 | 3 | 1 | 2 | 2 | 2 | 6 | | 7 | 1 | 3 | 2 | 1 | 4 | | 4 | 1 | 8 | |
| S metabolism | Sulfur cycling\|Sul de oxidation | | 1 | 5 | | 4 | 1 | | 6 | | | 1 | | | 1 | | | | 3 | 23 | | | | | | | | | 3 | | | | | | | | | | | | |
| S metabolism | Sulfur cycling\|Sul te reduction (Asr) | | | | | | | | | | | | | | | | | | | | | | | | | | | | | | | | | | | | | | | | |
| S metabolism | Sulfur cycling\|Sul te reduction (Dsr) | | | | | | | | | | | | | | | | | | | | | | | | | | | | | | | | | | | | | | | | |
| S metabolism | Sulfur cycling\|Sulfur oxidation | 6 | | 13 | 3 | 13 | 1 | 1 | | 3 | 4 | 8 | 4 | 1 | 13 | | 1 | 1 | 4 | 41 | 1 | 1 | | 3 | 2 | 2 | 2 | 2 | 6 | 1 | 7 | | 7 | 2 | 1 | 2 | 1 | 4 | 1 | 2 | 2 |
| S metabolism | Sulfur cycling\|Sulfur reduction | | | | | | | | | | | | | | | | | | | | | | | | | | | | | | | | | | | | | | | | |
| S metabolism | Sulfur cycling\|Thiosulfate disproportionation | | | | | | | | | | | | | | | | | | | | | | | | | | | | | | | | | | | | | | | | |
| S metabolism | Sulfur cycling\|Thiosulfate oxidation | 2 | | 1 | | | | | 6 | | | | | | | | | | | 26 | | | | | | | | | | | | | | | | | | | | | |
| Other metabolisms | Hydrogenases | 4 | 3 | 5 | 1 | 8 | 2 | | 6 | 5 | 2 | 7 | 3 | | 11 | 1 | 1 | 1 | 3 | 22 | | 1 | 1 | 3 | 2 | 2 | | 2 | 8 | | 3 | 1 | 5 | | | 6 | 1 | | | 7 | 1 |
| Other metabolisms | Urea utilization | 2 | | 5 | | | | | | | | | 1 | | | | | | | 24 | | | | | | 1 | 1 | | 2 | | | | | | | | | | | 1 | |
| Other metabolisms | Halogenated compound utilization | 4 | 1 | 3 | | 4 | | | | 2 | | 1 | | | | | | | | 25 | | | 1 | | | 2 | | 1 | | | | | | | | 1 | 1 | 1 | 1 | 6 | |
| Other metabolisms | Perchlorate reduction | 2 | | 2 | | 5 | 1 | | 3 | 2 | 1 | | 1 | | 1 | | | | 3 | 7 | 1 | | | 2 | | | | 2 | 1 | | 1 | | | | | | | 1 | | | |
| Other metabolisms | Chlorite reduction | 3 | | 1 | 1 | 3 | | | 5 | | | 4 | 1 | | | | | | | 8 | | | | | | 2 | 1 | | 2 | | | | | 2 | | | | | | | |
| Other metabolisms | As cycling\|Arsenate reduction | 10 | 1 | 12 | 3 | 34 | 7 | 1 | 5 | 6 | 2 | 7 | 3 | 1 | 18 | 1 | 1 | 3 | 2 | 45 | 1 | 1 | 1 | 3 | 6 | 2 | 2 | 1 | 6 | | 5 | | 4 | 2 | 1 | 8 | | 3 | 1 | 9 | 1 |
| Other metabolisms | As cycling\|Arsenite oxidation | | | 2 | 1 | | | | | | | | 1 | | | | | | 1 | 5 | | | | | | 1 | | | | | | | | | | | | | | | |
| Other metabolisms | Selenate reduction | 1 | 1 | | 2 | 13 | 2 | 1 | | 4 | | 2 | | | 10 | | | 1 | | 2 | | | | | | | 1 | | | | | | 2 | | 1 | 3 | | | | 4 | |
| Other metabolisms | Nitrile hydration | | | 1 | | | | | | | | | | | | | | | | 6 | | | 1 | | | 1 | | | | | | | | | | | | | | | |
| Other metabolisms | Metal reduction | 10 | | | 2 | 1 | 2 | | | | | 3 | 3 | | 3 | | | | 2 | 21 | | | 1 | 3 | | 2 | 2 | 1 | 2 | | | | | | 1 | | | 1 | 1 | | 2 |

**Supplementary Figure S6. Metabolic profile diagram of Rifle subsurface microbial community.** The dereplicated genomes from Rifle subsurface were assigned with genome taxonomy by GTDB-Tk, and they are clustered into 40 microbial groups. The metabolic functional traits were summarized and represented accordingly.
