## Supplementary Figure S7 for "METABOLIC: High-throughput profiling of microbial genomes for functional traits, biogeochemistry, and community-scale metabolic networks"

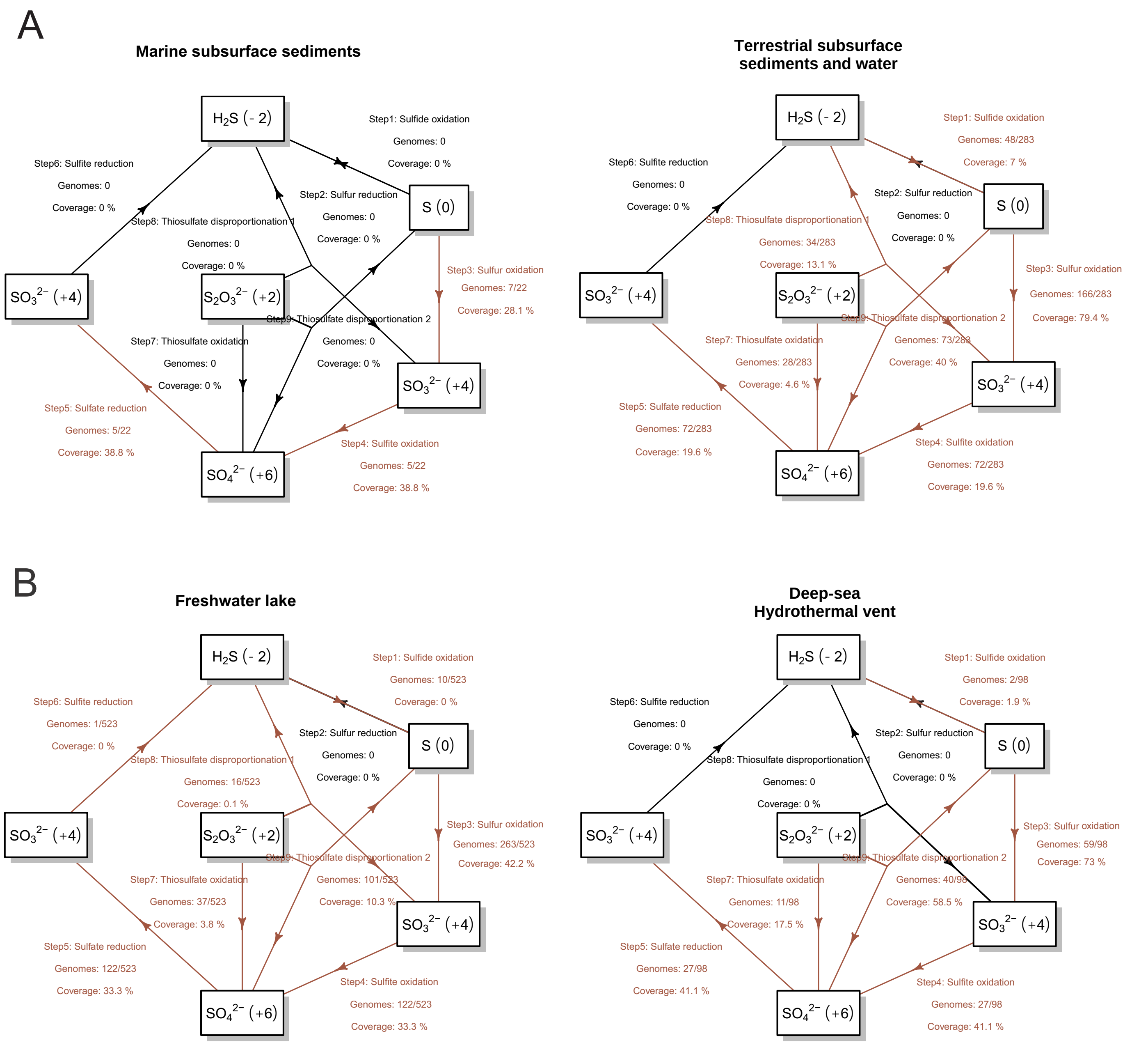


**Supplementary Figure S7. The sulfur related metabolism comparison at the community scale level.** (A) Comparison between Deep subsurface sediments (marine subsurface) and Rifle (terrestrial subsurface). (B) Comparison between Lake Tanganyika (freshwater) and Guaymas Basin (deep-sea plume). Each arrow represents a single transformation/step within a cycle. Indicated above the arrows in sequential order from top to bottom are: Step number and reaction, number of genomes that can conduct these reactions (and the total genome number of the community), metagenomic coverage expressed as a percentage of the community. The dark red labeled arrows and tags indicate the presence of the transformation/step within the community.
